# Extracellular Vesicle-Linked Vitamin B_12_ Acquisition via Novel Binding Proteins in *Bacteroides thetaiotaomicron*

**DOI:** 10.1101/2025.07.31.667871

**Authors:** Rokas Juodeikis, Robert Ulrich, Charlea Clarke, Michal Banasik, Evelyne Deery, Gerhard Saalbach, Bernhard Kräutler, Simon R. Carding, Michael A. Geeves, Richard W. Pickersgill, Martin J. Warren

## Abstract

The acquisition and utilization of vitamin B_12_ (cobalamin) are essential for the metabolic functions of many gut bacteria, including *Bacteroides thetaiotaomicron*, which relies on external sources of cobalamin due to its inability to synthesize it *de novo*. To scavenge cobalamin efficiently, *B. thetaiotaomicron* employs a sophisticated cobamide uptake system comprising multiple operons encoding outer membrane binding proteins and transporters. This study identifies and characterizes several novel cobalamin-binding proteins and elucidates their roles in cobamide uptake and delivery via bacterial extracellular vesicles (BEVs), highlighting their siderophore-like function for competitive nutrient acquisition.

We demonstrate that BtuJ1 and BtuJ2, members of the IPR027828 protein family, bind cobalamin with distinct structural and kinetic properties and are key components of BEVs responsible for cobamide capture and transfer to cells. Structural analyses reveal a conserved augmented β-jellyroll architecture in these proteins, with tyrosine residues playing a central role in stabilizing cobalamin binding. Comparative proteomics of BEVs and cells under cobamide starvation underscore the selective enrichment of BtuJ proteins in BEVs, suggesting a specialized mechanism for nutrient acquisition. Additionally, we identify another novel B_12_-binding protein, BtuK1.

We further establish BtuL as a critical player in early BEV release and propose a theoretical model in which BEVs function similarly to siderophores, scavenging cobalamin in the environment and delivering it to cells via specific receptors. This study provides new insights into the interplay between BEV-mediated transport, cobamide uptake, and the metabolic strategies employed by gut bacteria to thrive in nutrient-limited environments.

## Introduction

Nutrient acquisition is fundamental to bacterial metabolism, growth and adaptation to changing environments. In addition to conventional uptake mechanisms, bacteria produce extracellular vesicles (bacterial extracellular vesicles, BEVs) that transport not only enzymes and signaling molecules but also sequester essential nutrients, such as iron and cobamides (a corrinoid compounds containing a central cobalt ion within a corrin ring (1)), from competing organisms and deliver them back to the bacterial cell (2–4) (Figure 1A). The acquisition of the cobamide cobalamin (vitamin B_12_) via BEVs is particularly crucial for *Bacteroides thetaiotaomicron*, a prominent commensal organism in the human gut (3, 5–9). Since *B. thetaiotaomicron* lacks the ability to synthesize cobalamin *de novo*, a highly complex and energetically demanding process (10, 11), it relies on external sources of this vitamin to support essential metabolic functions, most notably the activity of its B_12_-dependent methionine synthase (MetH) enzyme (3, 12). To optimize cobamide scavenging, *B. thetaiotaomicron* employs a sophisticated cobamide uptake system composed of multiple transport loci encoding outer membrane cobamide-binding proteins and transporters, including several previously uncharacterized outer membrane lipoproteins (Figure 1B) (6).

**Figure 1.**
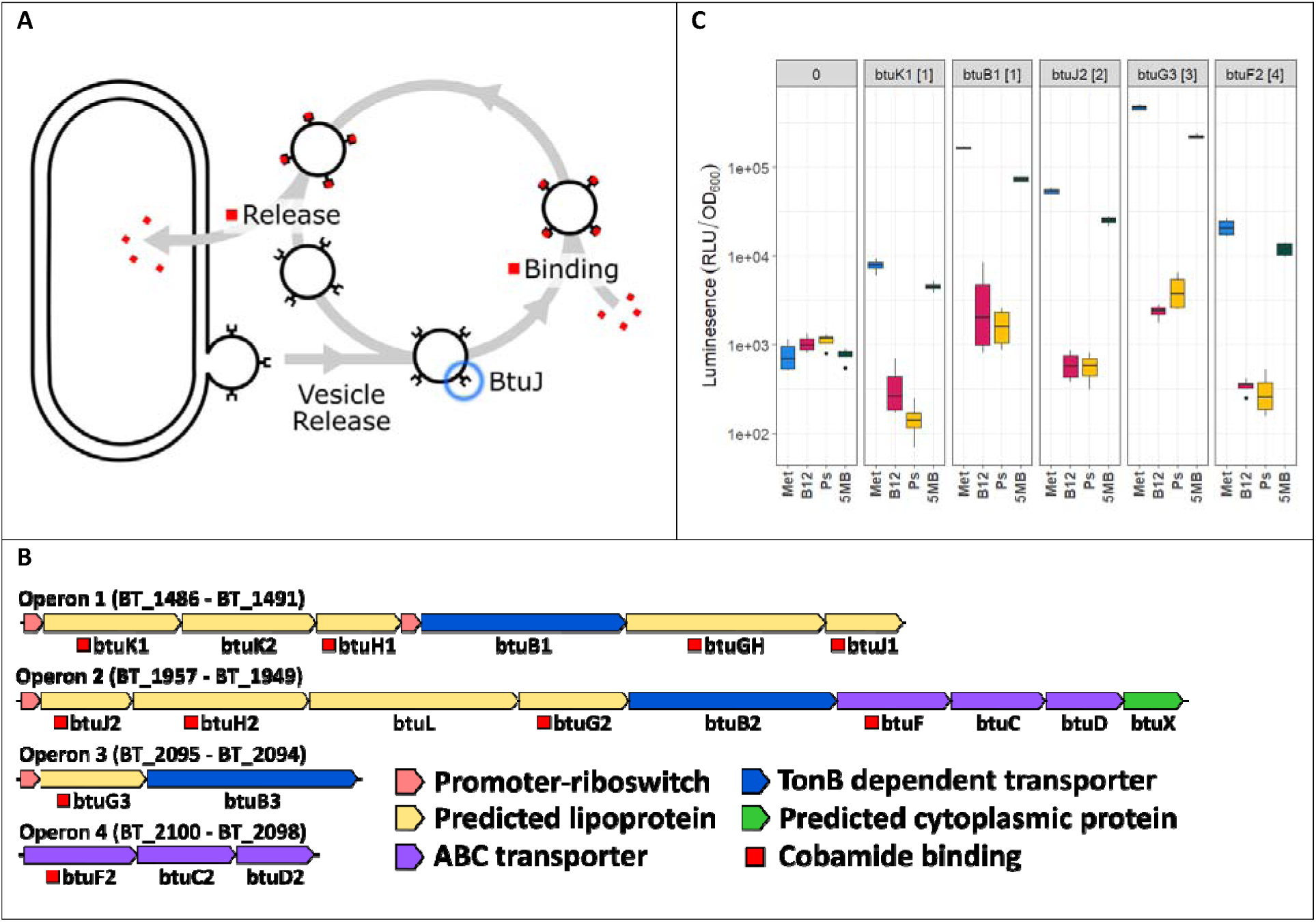
Cobamide uptake operons in *Bacteroides thetaiotaomicron* VPI-5482. A) Suggested model for nutrient uptake where proteins on extracellular vesicles act as siderophores, binding nutrients and delivering these to the cell in a receptor dependent manner. Vesicle enriched cobalamin binding protein discussed in this paper is highlighted in blue. B) Panel shows the genetic elements involved in cobamide uptake. The operons are regulated by a cobalamin riboswitch, apart from operon 4. Notably, operon 1 contains an additional internal promoter site. Cobamide binding annotations are based on this publication. C) Bar chart showing how identified promoters are regulated by different cobamides. The top label identifies different promoters, where 0 is a negative control, labels identify the location of the promoter, and the ribosome binding site used, and square brackets indicate the operon where the gene is located. The labels at the bottom indicate the test condition, where Met is the negative control with 400 µM L-methionine, and others contain different cobamides; B_12_ is cyanocobalamin, Ps is pseudocobalamin, 5MB is methylbenzimidazole cobamide. For all promoters, B12 and Ps show 100% downregulation, while 5MB shows around 50%. Boxplots are representative of four replicates

Recent studies have identified several high-affinity cobalamin-binding lipoproteins encoded within these loci, which are essential for nutrient acquisition and its subsequent transfer to the transporter (5, 6, 8, 9). The best characterized of these outer membrane binding proteins is BtuG, a protein crucial for the activity of the co-expressed TonB-dependent cobalamin transporter, BtuB. Interestingly, BtuG is not found in other previously characterized organisms that utilize BtuB (5, 6, 8, 9). Another strong cobalamin binder found within these operons, BtuH, has no clearly characterized biological function (8). The existence of this complex cobamide uptake system points to the highly competitive nature of its acquisition within the gut microbiota (6). The presence of multiple cobamide-binding proteins within this uptake system suggests a cooperative mechanism in which these proteins sequentially transfer the nutrient among themselves, facilitating its movement from the extracellular environment to specific transporters. This stepwise handoff likely enhances the efficiency of cobamide acquisition and ensures its effective internal accumulation, highlighting the intensely competitive nature of nutrient uptake within the gut microbiota (5).

*B. thetaiotaomicron* is highly adapted to the human gut, where competition for nutrients and space is intense. Its ability to produce BEVs enhances its adaptability, enabling dynamic interactions within this complex environment. In our previous work, we demonstrated that *B. thetaiotaomicron* BEVs bind and specifically deliver cobalamin, effectively sequestering it from competing organisms, such as *Salmonella* (3, 13). Moreover, we showed that these BEVs can enter host cells and efficiently deliver essential micronutrients to the host (3). The gastrointestinal tract contains a diverse array of cobamides, including cobalamin analogues in which the lower ligand, dimethylbenzimidazole, is replaced with adenine or modified adenine derivatives. Previous studies have demonstrated that *B. thetaiotaomicron* utilizes distinct BtuB transporters to selectively acquire these cobamides, conferring a competitive advantage in this nutrient-limited ecosystem (6). A deeper understanding of how cobamide-binding proteins interact with these structurally diverse molecules is crucial, particularly in elucidating why they exhibit such high binding affinity and how they are subsequently released to the appropriate transporters. Unraveling these mechanisms is key to understanding the efficiency and specificity of cobamide uptake. Such detailed characterization is essential for constructing a molecular framework to define the sequential events that drive competitive cobamide uptake and acquisition. Furthermore, it will shed light on how BEVs enhance this process, amplifying nutrient acquisition strategies and contributing to bacterial adaptation within the gut environment.

In this study, we further characterize the cobamide uptake system in *B. thetaiotaomicron.* We identify additional, previously uncharacterized cobamide-binding lipoproteins present in the cobamide uptake operons. Furthermore, we identify the specific proteins responsible for cobamide binding activity on BEVs and establish a link between BEV release and cobamide uptake. Our findings provide new insights into the mechanisms of cobamide acquisition and elucidate the roles of specific proteins in cobamide binding and delivery via BEVs.

## Results

### Operon analysis

Previous work identified and characterized four operons located across three genetic loci in *B. thetaiotaomicron* VPI-5482 that are involved in the uptake of a range of cobamides (Figure 1B) (6). Based on homology and gene number, we propose labelling the previously uncharacterized genes within these operons as *btuK1/2* (BT_1486/BT_1487), *btuJ1/2* (BT_1491/BT_1957), *btuL* (BT_1955), and *btuX* (BT_1949). Notably, with the exception of *btuX*, all of these are predicted to encode lipoproteins, as indicated by the presence of a signal motif identified using SignalP 6.0 (14). We further propose that BT_1486 has a misannotated start codon, suggesting methionine 36 is the correct start site. This is supported by the presence of a lipoprotein signal peptide starting at this sequence and the absence of the first 35 residues in the highly conserved BT_1487 gene. Details of our proposed operon annotations are available in Supplementary Dataset 1.

Operons 1, 2 and 3 are predicted to be regulated by cobalamin riboswitches, with operon 1 containing an additional riboswitch localized upstream of *btuB1* (Figure 1B) (6). These riboswitches are proposed to function by binding specific cobamides, inducing a structural rearrangement that leads to premature transcription termination and a consequent reduction in protein production (6, 15). In contrast, no riboswitch has been identified for operon 4.

To confirm the regulation of these operons, we generated reporter constructs using a single-copy integration vector targeting the promoter regions, including the site upstream of *btuB1*. Each construct incorporated the downstream ribosome binding site, which was used to name the constructs, to drive translation. Nanoluciferase was used as the reporter gene (16). The resulting reporter strains were analyzed under conditions with three different cobamides, using methionine (Met) as a control (Figure 1C). As expected, in the absence of cobamides and with methionine as the growth substrate, gene expression was induced. Among the operons, operon 3 exhibited the highest expression level, while the promoter upstream of operon 1 showed the lowest. Interestingly, the promoter upstream of *btuB1* within operon 1 demonstrated high levels of translation, confirming its role as an active promoter site.

In the presence of cobalamin (B_12_) and pseudocobalamin (Ps), all tested promoters exhibited near-complete repression of gene expression. However, exposure to 5-methylbenzimidazole cobamide (5MB) resulted in partial downregulation, reducing protein production to approximately 50%.

### Cobalamin binding lipoproteins

To gain deeper insight into of cobamide uptake in *B. thetaiotaomicron* and explore the role of the recently identified novel B_12_-binding proteins encoded within the four operons in salvaging cobamides from the growth media, we first sought to identify all the cobalamin-binding lipoproteins encoded within these regions. To achieve this aim, we recombinantly expressed each identified lipoprotein in *Escherichia coli*, replacing their N-terminal lipoprotein motif with a His-tag to facilitate purification. Proteins capable of binding cobalamin were identified based on their ability to retain the cofactor when bound to a nickel affinity column.

This approach identified a remarkable total of ten cobalamin-binding proteins encoded within the four operons (Figure 1B, red squares annotations). These include the previously identified BtuGH (formerly BtuG1), BtuG2/3 and BtuH1/2. Sequence analysis of BtuGH suggests that it represents a fusion of BtuG and BtuH domains. Upon dissecting this protein into its two component domains, we confirmed that both independently bind cobalamin. We also verified that the two predicted cobalamin ABC transporter periplasmic components (BtuF1 and BtuF2) are indeed cobalamin-binding proteins. Interestingly, BtuF2 contains a predicted lipoprotein sequence, though its functional significance was not explored in this study. Additionally, we identified three novel cobalamin binding proteins. Both BtuJ1 and BtuJ2 demonstrated B_12_ cobalamin-binding, whereas BtuK1, but not BtuK2, was also able to bind cobalamin. This was unexpected, given the high sequence similarity between BtuK1 and BtuK2. Finally, no cobalamin binding was observed for BtuL.

To understand better the roles of these cobalamin-binding proteins in sequestering exogenous cobamides and facilitating their transport to the cellular cytoplasm, we investigated their binding kinetics, given previous reports of femtomolar binding affinities. To achieve this, we employed stopped-flow spectroscopy and surface plasmon resonance (SPR) measurements to characterize their binding properties.

### Binding affinity measurements

Stopped flow measurements revealed that cobalamin binding to the Btu proteins caused a decrease in protein tryptophan fluorescence, by as much as 50% in some cases (Supplementary Figure 1A). A plot of the observed rate constant for binding (*k_obs_*) as a function of cobalamin concentration yielded straight lines, with the second-order binding rate constant (*k_on_*) determined from the slopes (Supplementary Figure 1B). The poorly defined intercepts suggest that *k_off_* is much smaller than the slowest *k_obs_* value predicting dissociation constants (*K_d_* = *k_off_*/*k_on_*) in the nanomolar range. In some cases, *k_off_* was determined by displacing cyanocobalamin from its complex with the Btu proteins using an excess of a second cobalamin-binding protein with distinct tryptophan fluorescence properties. For example, when cyanocobalamin bound to BtuJ1 was displaced by BtuJ2, the release rate constant (*k_off_*) was estimated at 0.03 s^-1^ (Supplementary Figure 1C), corresponding to an affinity (*K_d_*) of 0.16 nM. Cyanocobalamin binding to BtuJ1 exhibited the fastest association rate constant (*k_on_* = 0.2 nM^-1^s^-1^), consistent with diffusion-controlled ligand binding (typically 0.1 – 1 nM^-1^s^-1^) (17, 18). Other *k_on_* values ranged from 0.01 to 0.2 nM^-1^s^-1^, except for BtuG1, which showed a significantly slower *k_on_* of 0.0004 nM^-1^s^-1^. While stopped flow allows for the accurate determination of fast on-rates, most of the off rates from our stopped-flow data are poorly defined or not determined.

Surface Plasmon Resonance (SPR) was employed to resolve the challenging binding kinetics associated with the extremely fast and exceptionally slow rates observed by stopped flow with these proteins. Experiments were conducted using the Biacore T200 system, which offers high sensitivity, optimized fluidics, and advanced data analysis capabilities (Supplementary Figure 1D). Compared to the stopped-flow method, SPR yielded slower apparent on-rates, potentially due to the immobilization of proteins on the surface support, which could affect binding dynamics (Table 1). Nonetheless, SPR enabled accurate determination of off-rates, which were generally slower than those measured using the stopped-flow technique. SPR analysis estimated cyanocobalamin affinities ranging from 0.037 nM for BtuG1 to 11 nM for BtuH1. Given the methodological advantages of SPR in measuring dissociation events, these off-rates are likely to be more reliable than those obtained via stopped-flow. Integrating the faster on-rates from stopped-flow measurements with the slower off-rates from SPR analysis suggests that proteins such as BtuG2, BtuG3 and BtuK1 could possess binding affinities in the picomolar range. However, additional studies with more refined methodologies will be necessary to obtain definitive affinity measurements.

**Table 1:** Summary of kinetics parameters obtained from stopped flow and SPR. ND – not defined.

|  | Stopped Flow |  |  | Surface Plasmon Resonance |  |  |
| --- | --- | --- | --- | --- | --- | --- |
| Protein | $k_{on} (s^{-1}M^{-1})$ | $k_{off} (s^{-1})$ | $K_d (M)$ | $k_{on} (s^{-1}M^{-1})$ | $k_{off} (s^{-1})$ | $K_d (M)$ |
| BtuF | $8.8 \times 10^7 \pm 0.2$ | $3.3 \times 10^{-1} \pm 9.8$ | $3.8 \times 10^{-9} \pm 11.1$ | ND | ND | ND |
| BtuF2 | $6.20 \times 10^7 \pm 0.05$ | ND | ND | $6.60 \times 10^5 \pm 0.07$ | $5.7 \times 10^{-3} \pm 0.04$ | $8.6 \times 10^{-9} \pm 0.1$ |
| BtuG1 | $3.4 \times 10^5 \pm 0.2$ | $4.8 \times 10^{-3} \pm 4.2$ | $1.4 \times 10^{-8} \pm 1.2$ | $6.40 \times 10^5 \pm 0.03$ | $2.3 \times 10^{-5} \pm 0.2$ | $3.70 \times 10^{-11} \pm 0.01$ |
| BtuG2 | $6.2 \times 10^7 \pm 0.1$ | $9.3 \times 10^{-1} \pm 16$ | $1.5 \times 10^{-8} \pm 2.6$ | $5.20 \times 10^5 \pm 0.02$ | $2.2 \times 10^{-5} \pm 0.1$ | $4.2 \times 10^{-11} \pm 0.3$ |
| BtuG3 | $5.80 \times 10^7 \pm 0.08$ | ND | ND | $2.00 \times 10^5 \pm 0.01$ | $4.0 \times 10^{-5} \pm 0.1$ | $2.00 \times 10^{-10} \pm 0.05$ |
| BtuH1 | $9.5 \times 10^6 \pm 0.2$ | $9.0 \times 10^{-2} \pm 10$ | $9.5 \times 10^{-9} \pm 10.5$ | $5.10 \times 10^5 \pm 0.05$ | $5.60 \times 10^{-3} \pm 0.03$ | $1.1 \times 10^{-8} \pm 0.01$ |
| BtuH2 | $8.9 \times 10^6 \pm 0.2$ | $2.0 \times 10^{-1} \pm 7.2$ | $2.3 \times 10^{-8} \pm 8.1$ | ND | ND | ND |
| BtuJ1 | $2.10 \times 10^8 \pm 0.07$ | $3.2 \times 10^{-2} \pm 0.2$ | $1.5 \times 10^{-10} \pm 0.1$ | $1.3 \times 10^7 \pm 0.01$ | $1.10 \times 10^{-2} \pm 0.01$ | $8.46 \times 10^{-10} \pm 0.10$ |
| BtuJ2 | $2.20 \times 10^8 \pm 0.03$ | ND | ND | ND | ND | ND |
| BtuK1 | $1.80 \times 10^7 \pm 0.02$ | $2.9 \times 10^{-1} \pm 5.0$ | $1.6 \times 10^{-8} \pm 2.8$ | $1.80 \times 10^5 \pm 0.01$ | $1.60 \times 10^{-5} \pm 0.08$ | $8.9 \times 10^{-11} \pm 0.5$ |

### The structures of BtuJ1 and BtuJ2 with bound cyanocobalamin

BtuJ1 and BtuJ2 share 23% amino acid sequence identity and belong to the IPR027828 protein family. Recombinant BtuJ1, expressed without its first 26-residue lipidation signal sequence, was crystallized in complex with cyanocobalamin. The structure was solved using molecular replacement with the AlphaFold Molecular Replacement workflow in the CCP4i program suite (see Materials and Methods for details). The crystals belong to space group C2_1_ and contain two BtuJ1/cyanocobalamin complexes in the asymmetric unit. The structure was refined to 1.6 Å resolution with good stereochemistry, achieving R_work_ and R_free_ values of 0.191 and 0.216, respectively (data collection and refinement statistics are summarized in Supplementary Table 1). The two complexes in the asymmetric unit are structurally similar, with an RMSD of 0.095 Å across 199 equivalenced C1 atoms.

The structure of BtuJ2 was determined using the same workflow as for BtuJ1. BtuJ2 crystals belong to space-group P2_1_2_1_2 and contain two molecules in the asymmetric unit. The structure was solved to 2.7 Å resolution, with reasonable crystallographic residuals and stereochemistry (Supplementary Table 1). The two copies in the asymmetric unit of BtuJ2 exhibit an RMSD of 0.127 Å across 211 equivalenced C1 atoms.

BtuJ1 and BtuJ2 share a core jelly-roll β-barrel structure, where the polypeptide chain forms eight β-strands organized into two four-stranded antiparallel β-sheets. This core structure is augmented by three additional antiparallel β-strands at the N-terminal end of the jellyroll and an α-helix located between strands D and E in the traditional jellyroll nomenclature (BIDG and CHEF, Figure 2A, B). When comparing BtuJ1 and BtuJ2, they exhibit a common core architecture (RMSD of 4.0 Å across 110 equivalent C1 atoms) but display significant differences in their loop regions. Notably, the HI loop, which contains a conserved tyrosine residue at its terminus near the start of strand I, is the most similar between the two proteins. This conserved loop is likely the key region involved in cobalamin-binding.

**Figure 2.**
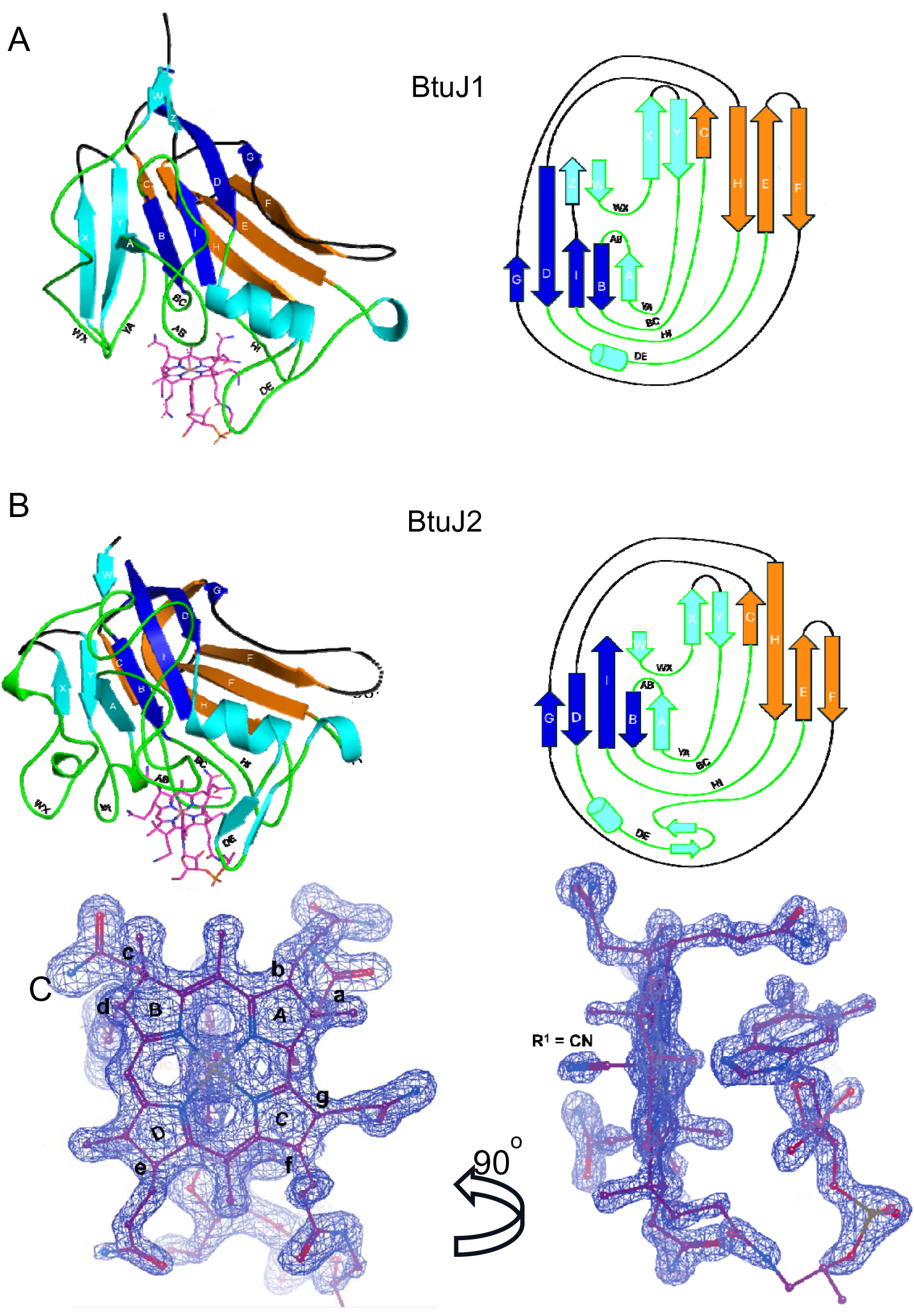
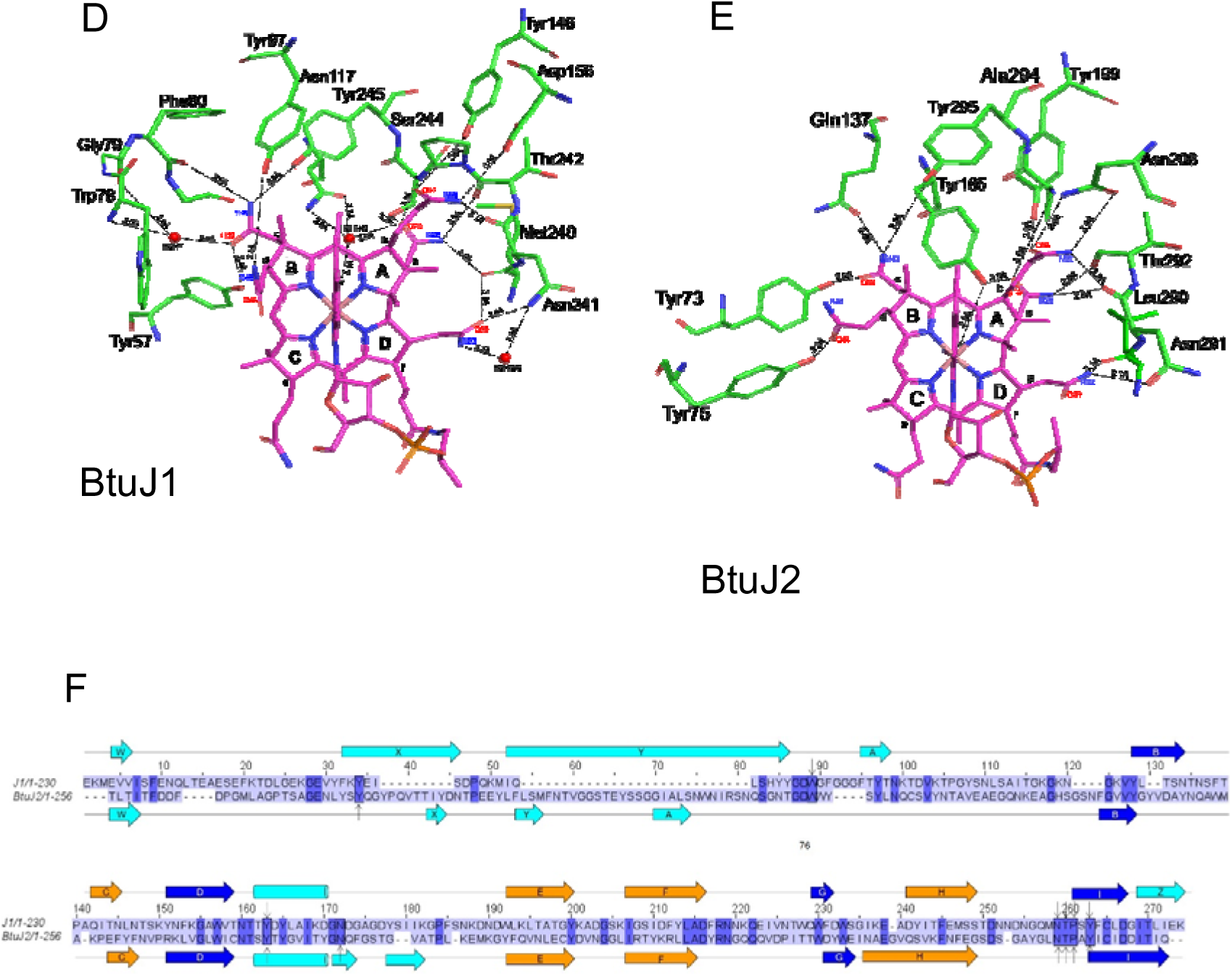
Crystal structure of BtuJ1 and BtuJ2 with bound cobalamin. The structure of BtuJ1 and BtuKJ2 with cobalamin bound (PDB: 9FCT and 9I2L). A. The augmented β-jellyroll architecture of BtuJ1 with additional β-strands (and helix) in light blue. Four loops (AB, BC, DE, HI) are directly involved in binding to cobalamin. The topology of BtuJ1 is shown on the right. B. The architecture of BtuJ2 and its topology. There is a lot of variation in β-strand length and loop structures compared to BtuJ1. C. Refined electron density for cobalamin bound to chain A of BtuJ1 in the crystal structure. Two orthogonal views are shown. D, E. Cartoons showing the hydrogen-binding residues of BtuJ1 and BtuJ2, respectively. There is an approximately equal mix of mainchain and sidechain hydrogen bonds. F. The sequence alignment of ButJ1 and BtuJ2 which have 23% sequence identity (conserved residues in blue block). Residues in the black boxes are conserved residues found hydrogen bonding to B12 in the crystal structures of J1 and J2. The arrow indicates the protein that the H bonding residue belongs to. The conserved hydrogen bonding residues from both are Tyr146 and Tyr199, Asn291 and Asn241, Thr242 and Thr292, Tyr245 and Tyr295. Arrows are β-strands and rectangles are α-helices and are labelled.

Cyanocobalamin is well-resolved in both crystal structures, with evidence of binding shown for BtuJ1 in Figure 2C. The cobalamin-binding pocket is formed by four loops from the jellyroll structure, loops: AB, BC, HI and DE and two additional loops associated with the ancillary β-strands; AB and ZA (Figure 2A, B). Within the pocket, the adenosyl group of the lower ligand faces outward, extending into the solution. Both binding pockets have sufficient space to accommodate the bulkier adenosyl-group of Ado-Cbl, and binding would displace water molecules, suggesting an entropic contribution to tight binding of Ado-Cbl.

The binding is predominantly characterized by shape complementarity between the pocket and the corrin ring, complemented by hydrogen-bonding. These interactions involve sidechain polar groups as well as mainchain carbonyls. When the cobalamin molecules are superimposed, the core architecture of BtuJ1 and BtuJ2 aligns well, highlighting similarities in the binding residues (Figure 2D, E, F).

A notable feature in both proteins is a halo of tyrosine residues that contribute hydrophobic interactions as well as hydrogen bonding to the corrin ring’s side chains. Conserved tyrosine residues in analogous positions include (BtuJ2 residue number in parentheses): Tyr 97 (134) on AB loop, Tyr 146 (199) at N-terminus of the helix within the DE loop (hydrogen bonding the *b* sidechain of the corrin ring), and Tyr 245 (295) at the end of the HI loop/start of strand I (hydrogen bonding to the *c* sidechain of the corrin ring).

An interesting distinction is observed in BtuJ1, where Trp 78 is involved in interactions with the dimethylbenzimidazole group of cobalamin. In BtuJ2, this tryptophan is replaced by a pair of tyrosine residues, resulting in a modification of the interaction with the cobalamin side chain (*d*), which was sandwiched between the dimethylbenzimidazole group and Trp 78 in BtuJ1. Overall, while the shape, hydrophobicity and hydrogen bonding characteristics of the binding site are retained in both structures, the specific residues contributing to these interactions differ, reflecting subtle variations in binding mechanisms between BtuJ1 and BtuJ2.

The resemblance between the six loops of the immunoglobulin fold and those of this augmented β-jellyroll has not gone unnoticed. This similarity raises intriguing possibilities for exploring the ligand-binding diversity within this jellyroll protein family.

### IPR027828 protein family analysis

The major binding motif (MNTPSY in *B. thetaiotaomicron*) includes the highly conserved Tyr245, a hallmark across the IPR027828 protein family (Domain of Unknown Function DUF4465). Proline and threonine residues occur frequently within this motif, and the tyrosine is universally present at the end of loop 6 (L6). This loop is a distinct and recognizable feature in all sequences for which AlphaFold models are available. Many of the interactions with the cobalamin ring are mediated by main-chain carbonyl oxygens, suggesting that this cobalamin binding site is likely conserved across the IPR027828 protein domain family. To understand further the localization of IPR027828 family members, we analyzed their targeting signals using SignalP 6.0 (14) and examined co-occurrences with InterPro domains associated with protein targeting. Out of 824 non-redundant sequences investigated, 38% contained InterPro annotation for a C-terminal extracellular targeting motif (229/824 IPR026444, 88/824 IPR013424), 33% contained a lipoprotein targeting sequence (275/824) while 21% contained a periplasmic targeting sequence (170/824). Notably, none of these targeting motifs co-occurred within individual sequences. Based on these observations, we propose that the IPR027828 family primarily consists of extracellular cobalamin-binding proteins, reflecting their likely role in nutrient acquisition and transport.

### Comparative cell and BEV proteomics under cobamide starvation

In our previous work, we demonstrated that *B. thetaiotaomicron* BEVs can bind and deliver a range of cobamides (3). To determine which of the identified cobalamin-binding proteins are involved in this BEV-mediated process, we conducted comparative proteomic analyses of cells and BEVs produced in cobamide-containing and cobamide-free media (Table 2; Supplementary Datasets 2 and 3). As expected, due to the regulation of these operons by cobalamin riboswitches, most of these proteins were significantly upregulated in both cells and BEVs when cobalamin was absent from the media. Similar regulatory patterns have been reported in recent studies (5). Notably, the protein upregulation profiles differed between cells and BEVs. The highest relative enrichment in BEVs was observed for BtuK2, BtuH1, BtuJ1 and BtuL, suggesting selective incorporation of these proteins into BEVs. Interestingly, BtuJ2 was undetectable in cells but showed significant upregulation in BEVs, highlighting its potential specialized role in vesicle-mediated cobamide transport.

**Table 2.** Proportional upregulation of cobamide uptake operon proteins in cells and BEVs produced in B12 free media. N/A = No signal detected. Statistical significance: *** p < 0.001, ** p < 0.01, * p < 0.05, N/S p > 0.05. SignalP values: LIPO = Lipoprotein (Red), SP = Signal Peptide (Light Blue), OTHER = Not detected (Pink). Gene colors highlight operons: 1 = Orange, 2 = Blue, 3 = Green, 4 = Yellow.

| Protein | Gene | SignalP Type | Cell (Met/B12) | BEV (Met/B12) |
| --- | --- | --- | --- | --- |
| BtuK1 | BT_1486 | LIPO | 81.1 (***) | 291.9 (***) |
| BtuK2 | BT_1487 | LIPO | 104.5 (***) | 1000 (***) |
| BtuH1 | BT_1488 | LIPO | 123.5 (***) | 1000 (***) |
| BtuB1 | BT_1489 | SP | 12.2 (***) | 79.7 (***) |
| BtuGH | BT_1490 | LIPO | 12.5 (***) | 21.8 (*) |
| BtuJ1 | BT_1491 | LIPO | 31.6 (***) | 471.3 (***) |
| BtuJ2 | BT_1957 | LIPO | N/A | 27.3 (***) |
| BtuH2 | BT_1956 | LIPO | 102.4 (***) | 259.9 (***) |
| BtuL | BT_1955 | LIPO | 99.6 (***) | 1000 (***) |
| BtuG2 | BT_1954 | LIPO | 74 (***) | 158.8 (***) |
| BtuB2 | BT_1953 | SP | 22 (***) | 45.4 (**) |
| BtuF | BT_1952 | LIPO | 208.8 (***) | 112.2 (***) |
| BtuC | BT_1951 | OTHER | N/A | N/A |
| BtuD | BT_1950 | OTHER | 13.9 (***) | 8.1 (N/S) |
| BtuX | BT_1949 | OTHER | 5.8 (***) | N/A |
| BtuB3 | BT_2095 | OTHER | 4.8 (**) | 47.3 (**) |
| BtuG3 | BT_2094 | LIPO | 47.9 (***) | 118.1 (***) |
| BtuF2 | BT_2100 | LIPO | 0.7 (N/S) | 0.5 (N/S) |
| BtuC2 | BT_2099 | OTHER | N/A | N/A |
| BtuD2 | BT_2098 | OTHER | 0.9 (N/S) | N/A |

In our recent study, we characterized the dynamics of BEV production by *B. thetaiotaomicron*, demonstrating that BEVs released early during growth are enriched with specifically targeted lipoproteins, whereas those released later contain whole cellular components, likely as a result of cell lysis (13). Using principal component analysis of the temporal proteomic data, we identified BtuK1, BtuK2, BtuJ1 and BtuL as major contributors to the BEV protein composition at distinct time points (Supplementary Figure 2). This finding suggests that these proteins are selectively incorporated into specialized, non-lytic BEVs during the early growth phase.

### BtuJ delivers cobalamin via BEVs

To determine whether any of these proteins contribute to cobalamin-binding activity in BEVs, we generated strains specifically expressing *btuK1, btuK2, btuJ1, btuJ2, btuH1, btuH2* or *btuL* in a mutant background lacking all cobamide uptake operons. BEVs purified from these strains were saturated with cobalamin, thoroughly washed, and provided to wild-type *B. thetaiotaomicron* cells as the sole source of cobalamin. The concentration of BEVs used was equivalent to that present in the late phase culture. Notably, only BEVs containing BtuJ1 or BtuJ2 served as an effective source for cobalamin (Figure 3A). We also tested BEVs derived from null mutant strains expressing *btuG* or *btuB* genes, which showed no significant binding activity (Supplementary Figure 3). These findings suggest that BtuJ1 and BtuJ2 are critical for cobalamin uptake via BEVs.

**Figure 3.**
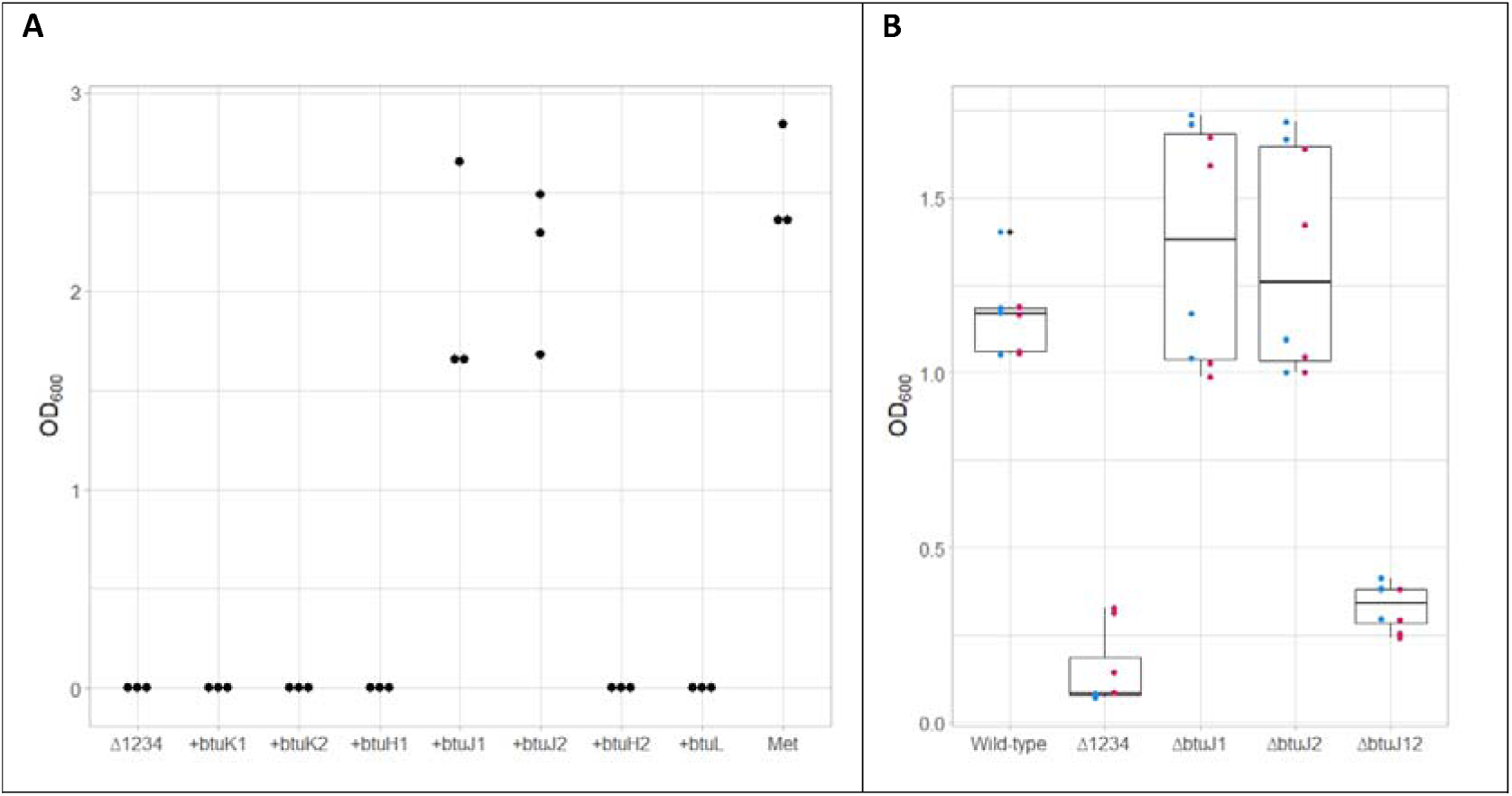
BtuJ1 and BtuJ2 are responsible for cobalamin delivery from *B. thetaiotaomicron* bacterial extracellular vesicles. BEVs from a range of generated *B. thetaiotaomicron* strains were used as the only source of cobalamin for wild-type *B. thetaiotaomicron*. A. Bioassay showing that either BtuJ1 or BtuJ2 is sufficient for cobalamin binding and delivery activity of BEVs. Δ1234 - *B. thetaiotaomicron* strain lacking all cobamide uptake genes; [+ gene] strains. Individual genes were reintroduced into *Δ1234* genetic background; Met is the positive control containing 400 µM L-methionine. The bioassay was in triplicate with one biological BEV replicate. Results for additional genes available as Supplementary figure 3. B. Growth recovery bioassay using cobalamin bound BEVs showing that either btuJ1 or btuJ2 are required for cobalamin binding and delivery activity. Wild type used as the positive control; Δ1234 used as the negative control; *ΔbtuJ1, ΔbtuJ2, ΔbtuJ12* are specific btuJ1, btuJ2 or double btuJ1 and btu2 knockout strains, respectively. The bioassay had 4 biological replicates of 2 biological replicates of BEV preparations indicated by colored dots with boxplots showed in black.

To validate this observation, we constructed *ΔbtuJ1*, *ΔbtuJ2* and *ΔbtuJ1/2* knockout strains of *B. thetaiotaomicron*. As before, BEVs from these strains were saturated with cobalamin, washed, and used at biologically relevant levels to rescue growth in cobamide-free media (Figure 3B). BEV concentrations were equivalent to those found in late-phase cultures. While BEVs from individual knockout strains (Δ*btuJ1* or Δ*btuJ2*) showed no significant effect on cobamide uptake, BEVs from the double knockout strain (Δ*btuJ1*/*J2*) exhibited a nearly complete loss of cobalamin-binding activity. These results strongly support the hypothesis that BtuJ1 and BtuJ2 are the primary proteins responsible for cobamide binding in BEVs and their subsequent release to cells.

### Cellular elements required for BEV-cobalamin uptake

Significant progress has been made in understanding the minimal cellular requirements for cobalamin uptake; however, certain specific combinations of gene knockouts have not yet been explored (6). Additionally, the role of BEV-associated cobalamin in this process remains unaddressed. To extend these previous findings and determine the cellular components necessary for utilizing BEV-bound cobalamin, we generated a series of novel knockout strains in the wild-type *B. thetaiotaomicron* background. These included strains with targeted operon deletions, such as Δ12, Δ23, Δ134 and Δ1234, as well as a complex knockout strain (Δ13.J2.H2.L) lacking operons 1 and 3 along with *btuJ2*, *btuH2*, and *btuL,* while retaining the promoter region of operon 2.

Growth experiments for these strains were conducted across a range of cobalamin concentrations, including conditions where BEV-associated cobalamin was the sole source (Figure 4A). Notably, differences in growth were observed at low cobalamin concentrations that were not apparent at higher levels. Interestingly, the Δ13.J2.H2.L mutant strain outperformed the wild-type strain under low cobalamin concentrations. A similar trend was observed when BEV-associated cobalamin served as the only source of cobalamin (Figure 4B).

**Figure 4.**
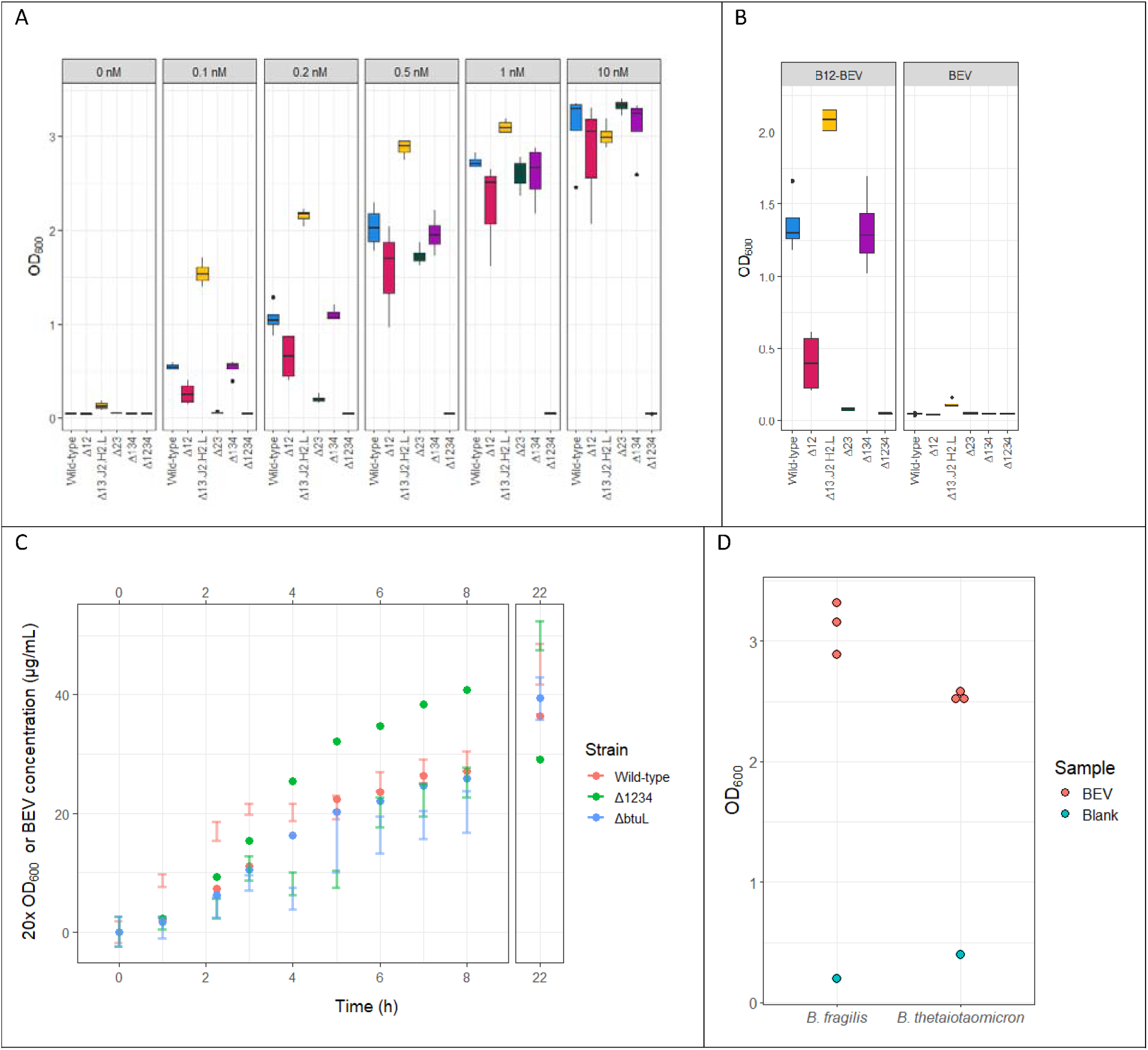
*Bacteroides* cobalamin uptake operon mutant analysis. A. Different *B. thetaiotaomicron* strain growth in the presence of varying cobalamin concentrations. Boxplots show 4 biological replicates. B. Different *B. thetaiotaomicron* strain growth utilizing BEV bound cobalamin as the only source (B12-BEV) and a cobalamin-free BEV control (BEV). Boxplots show 4 biological replicates of one BEV preparation. C. BEV release over time by different *B. thetaiotaomicron* mutant strains highlighting the role of BtuL. Dots show representative average 20*OD_600_ value change from inoculation; bars show BEV concentration in the supernatant as one standard deviation in 4 biological replicates. Note the difference in BEV release between 1 and 5 hours of culture. D. *B. fragilis* utilization of cobalamin bound to *B. thetaiotaomicron* BEVs.

Based on the enhanced performance of the Δ13.J2.H2.L strain, the significant upregulation of BtuL in BEVs, and the absence of detectable cobalamin binding by BtuL, we hypothesized that BtuL might play a role in promoting BEV release. To test this, we measured BEV release over time in wild type, Δ1234, and Δ*btuL B. thetaiotaomicron* strains (Figure 4C). Notably, both mutant strains produced fewer BEVs during the early growth phase compared to the wild-type strain. Interestingly, this difference was no longer observed in the later stages of growth.

In our previous work, we demonstrated that BEV-associated cobalamin is not bioavailable to an auxotrophic *Salmonella* strain (3). To investigate whether BEV-bound cobalamin is accessible to closely related *Bacteroides* species, we tested its bioavailability in *Bacteroides fragilis* NCTC 9343, a strain that lacks most of the cobalamin uptake genes present in *B. thetaiotaomicron* (Supplementary Figure 4). Interestingly, this *B. fragilis* strain was able to utilize BEV-bound cobalamin (Figure 4D). However, BEVs produced by *B. fragilis* did not exhibit cobalamin-binding activity. This finding further supports the role of BtuJ as the key cobalamin-binding protein on BEVs, as BtuJ is absent in this particular *B. fragilis* strain.

## Discussion

In this study, we expand on previous work identifying two novel cobalamin-binding proteins, BtuK1 and BtuJ, in the model gut commensal bacterium *B. thetaiotaomicron*. We provide detailed structural and binding kinetic analyses of BtuJ1 and its homologue, BtuJ2. Additionally, we demonstrate the critical roles of BtuJ1 and BtuJ2 in cobalamin binding on BEVs and elucidate the role of BtuL, a protein encoded in the cobalamin uptake operon 2, in promoting BEV release.

Bioinformatic analysis of the four cobalamin uptake operons in *B. thetaiotaomicron* VPI-5482 revealed the presence of an internal promoter in operon 1 and the absence of a riboswitch in operon 4. Recombinant promoter-reporter assays demonstrated that this internal promoter is not only active but also approximately an order of magnitude stronger than the primary promoter at the start of the operon. This suggests relatively low expression of *btuK1, btuK2* and *btuH1*. Despite the lack of an identifiable riboswitch in operon 4, its gene expression was regulated similarly to the other cobalamin uptake operons, indicating the presence of an alternative regulatory element potentially functioning in place of a riboswitch. Additionally, we propose a novel and comprehensive naming convention for all the identified genes involved in cobamide uptake to standardize future references and analyses.

This study provides further evidence of *B. thetaiotaomicron* BEVs playing a critical role in cobamide capture by identifying BtuJ proteins as cobamide-binding components localized to BEVs and implicating BtuL as a key player in early-stage BEV release. Notably, our experiments focused specifically on BEV proteins involved in both cobamide binding and release; it remains possible that other cobalamin-binding proteins are present on BEVs either in lower abundance or without the ability to release bound cobalamin. For example, while we observed significant enrichment of BtuK2 and BtuH1, their precise roles remain to be determined.

In our previous work, we characterized different types of BEVs, finding that BEVs released before mid-log phase are enriched in lipoproteins, including several Btu proteins (13). When we examined temporal BEV release in a null cobamide uptake operon mutant and a Δ*btuL* strain, we observed a reduction in these early-phase BEVs. This supports the hypothesis that BEVs released during this stage of growth are specialized for metabolic functions. Interestingly, the null mutant strain exhibited better growth than the wild-type strain, suggesting that BEV release and the expression of cobamide uptake operons impose a significant metabolic burden on the cell. While the experimental conditions used in this study do not fully replicate the natural gut environment, these findings underscore the substantial metabolic investment made by cells to facilitate cobamide capture.

BtuJ1 and BtuJ2 share a common augmented β-jelly-roll architecture and bind cobalamin at the same site and in the same orientation. However, this binding occurs with limited conservation of binding residues. The greatest conservation is observed in the HI loop, which features a conserved tyrosine residue. Tyrosine residues play a central role in cobalamin binding, forming a “halo” that contributes hydrophobic interactions and stabilizes the corrin side chains. The diversity observed in the binding loops and residue positions suggests that cobalamin binding is a predominant function of the DUF4465 protein superfamily, despite variations in the specific molecular interactions involved. The tight binding of cobalamin to proteins such as BtuJ1, J2 and K1 also holds potential for biotechnological applications.

We propose a theoretical model in which BEVs function similarly to siderophores, being released from cells to scavenge cobalamin in the environment by binding it tightly. Once bound, the vitamin is only made available to cells equipped with the appropriate receptor (Figure 1B), with its release likely facilitated by other proteins. From a structural perspective, BtuJ and BtuG bind cobamide with the lower ligand loop exposed, facing outward from the binding pocket, whereas BtuH binds on the opposite side, with the lower ligand loop-oriented inwards (5, 8). This arrangement supports a mechanism in which cobalamin captured by BtuB on BEVs is transferred to BtuH on the cell surface, and subsequently onto BtuBG complex. Further evidence for this mechanism comes from the presence of a BtuGH fusion protein in operon 1. This model would allow for a reduced number of BtuBG complexes on the cell surface if BtuH is present to mediate cobamide capture.

## Methods

### Bacterial culture conditions

*E. coli* was grown in lysogeny broth (LB) containing 10g/L Tryptone; 5g/L Yeast Extract; NaCl 10 g/L at 37 °C at 200 rpm when grown in liquid culture. *B. thetaiotaomicron* was grown in an anaerobic cabinet (10% H2, 5% CO2, 85% N2) at 37°C with no agitation. Brain Heart Infusion supplemented with 4-μM hemin (BHIH) or Bacteroides Defined Media r6 (BDMr6) consisting of 100 mM potassium phosphate pH 7.8 (KH_2_PO_4_ 9.2 mM; K_2_HPO_4_ 90.8 mM), 15 mM NaCl, 8.5 mM (NH_4_)_2_SO_4_; 30 mM glucose; 100 µM MgCl_2_; 50 µM CaCl_2_; 2 mM L-Cysteine hydrochloride; 10 μM FeSO_4_; 400 µM L-methionine (or specified concentration of L-methionine/cobamide); 50 nM Protoporphyrin IX (prepared in 50% ethanol; 50 mM NaOH) was used to culture cells. All components purchased from Merck unless otherwise specified.

### Molecular biology

*E. coli* DH5α (NEB) used to construct expression plasmids; *E. coli* PIR1^+^ (ThermoFischer Scientific) used for integration and knockout vectors. Chemically competent *E. coli* cells were used with transformations carried out using heat-shock method; *B. thetaiotaomicron* transformations carried out via conjugation using relevant *E. coli* donor strains as previously described (19). All primers and plasmids used are outlined in Supplementary Table 2 and Supplementary Table 3; all plasmid sequence maps are also available as Supplementary Dataset 1. Plasmid sequences were confirmed using sanger sequencing (Genewiz).

To generate the cobalamin promoter *B. thetaiotaomicron* reporter construct, a synthetic DNA fragment (Integrated DNA Technologies) containing a promoter (P.Bth_BT1830s); ribosome binding site (RBS.Bth_RBS7) and a codon optimized Nanoluciferase; flanked by terminators (T.BBa_B1001; T.BBa_B1007) was inserted into a single copy integration vector pNBU2_erm-TetR-P1T_DP-GH023 *AflII*/*BamHI* site yielding pIBATH.56 (20, 21). *BglII*/*NdeI* sites were encoded in the DNA fragment to allow for the replacement of promoter and RBS site. This site was used to insert DNA fragments corresponding to the promoter and RBS sites generated by PCR (Phusion; NEB) using outlined primers (Supplementary Table 2 and 3; Supplementary Dataset 1).

To generate plasmids for protein expression in *E. coli,* the soluble parts of the genes of interest, lacking the predicted lipoprotein motif as identified using SignalP 6.0 (11), were amplified via PCR using primers containing restriction site overhangs outlined in Supplementary Table 2. The amplified fragments were inserted into a modified pET14b vector (novel *SpeI* site between the *NdeI* and terminator) *NdeI/SpeI* site resulting in a construct encoding for an N-terminus hexahistidine tagged protein produced via the T7 promoter (Supplementary Table 2; Supplementary Dataset 1).

To generate single gene expression plasmids for *B. thetaiotaomicron*, pGH117 was used as the vector (19). The *Pci*/*BamHI* cloning site was used to introduce a synthetic promoter (P.Bth_BT1830s) and ribosome binding site (RBS.Bth_RBS7) followed by an *NdeI*/*SpeI* cloning site containing an ORF (not relevant to this study) with artificial bidirectional terminators (BBa_B1001; BBa_B1007) flanking the operon yielding pBATH.03 (21). PCR was used to amplify genes of interest using *B. thetaiotaomicron* VPI-5482 DNA and primers (Integrated DNA Technologies) designed with *NdeI/SpeI* flanking restriction sites, which were used to insert the generated fragments into the pBATH.03 vector *NdeI/SpeI* site (Supplementary Table 2 and 3; Supplementary Dataset 1). The generated plasmids were used to transform *B. thetaiotaomicron* strain lacking all three cobalamin uptake operons (*Δlocus1 Δlocus2 Δlocus3 Δtdk* strain) (6).

*B. thetaiotaomicron* VPI-5482 mutant strains were generated using a previously reported method (22). Plasmid for the crossover (Supplementary Table 3; Supplementary Dataset 1) were generated via insertion of a fragment generated by overlap extension PCR. Primers corresponding to downstream (Down), or upstream (Up) fragments were used to amplify DNA upstream and downstream of the target DNA (Supplementary Table 2; Supplementary Dataset 1). These fragments were combined using overlap extension PCR. Primers contained engineered restriction enzyme sites for *BamHI* (Up) and *PstI* (Down), which were used to insert the fragment into pLGB13 (22). Post conjugation, individual recovered colonies were grown overnight in BHIH containing gentamicin (200 µg/mL) and erythromycin (25 µg/mL) and streaked out on BHIH agar plates containing gentamicin (200 µg/mL) and anhydrotetracycline (100 ng/ml). Individual colonies were then tested using PCR for the loss of the target DNA segment. To generate multiple edit strains, this process was repeated.

### Promoter characterization

*B. thetaiotaomicron* VPI-5482 cobalamin promoter reporter strains were generated via transformation with the generated reporter plasmids (pIBATH.104-108; Supplementary Table 2; Supplementary Dataset 1). Two colonies for each transformation were selected and cultured in duplicate overnight in 5 mL BDMr6. 10 µL was used to inoculate 200 µL BDMr6 containing either 200 µM L-methionine; 500 pM cyanocobalamin (vitamin B_12_); 500 pM pseudocobalamin (Ps); or 500 pM methylbenzimidazole cobamide (5MB) in a flat-bottom 96 well plate (bio-one cellstar; Greiner), sealed with adhesive gas permeable seals to limit evaporation, overnight. Nanoluciferase signal was quantified using Nano-Glo® Luciferase Assay System (Promega) and adjusted to cell density as estimated by optical density at 600 nm (OD_600_). Readings carried out using CLARIOstar Plate Reader (BMG Labtech) using standard instrument settings.

### Protein expression

*E. coli* BL21(DE3) cells were used for protein production. Transformants selected using 100 µg/mL ampicillin. Single colonies were used to inoculate 50 mL LB (100 µg/mL ampicillin) and cultured overnight at 37°C. 8 mL of the overnight culture was transferred to 800 mL of fresh media and cultivated until the culture reached the mid-log phase, measured with an OD_600_ value of 0.6. At this point, the culture was cooled down and induced with IPTG (1 mM). The induced culture was incubated at 18°C overnight at 200 rpm. Bacteria were collected by centrifugation, and the obtained pellets were kept frozen at -20 °C until required.

### Protein purification

Bacterial pellet resuspended in 100 ml of binding buffer (20 mM Tris-HCl, pH 7.5, 500 mM NaCl, 30 mM imidazole, 1 mM TCEP), supplemented with complete protease inhibitor cocktail tablets (Roche), and sonicated on ice. The obtained lysates were clarified by centrifugation (16,000 x g, 30 min, 4°C), followed by ultracentrifugation (80,000 x g, 30 min, 4°C), and supplemented with 1 mL of cyanocobalamin stock solution (10 mg/mL). The resulting supernatant was passed five times through a pre-packed HisTrap HP column containing 5 mL of chelating Sepharose (Cytiva). Subsequently, the column was washed with 100 ml binding buffer, and the target protein was eluted in 5 ml fractions with elution buffer (20 mM Tris-HCl, pH 7.5, 500 mM NaCl, 500 mM imidazole, 1 mM TCEP). Following protein elution, the elution fractions containing the highest amount of total protein (estimated by absorbance at 280 nm) were combined, concentrated by centrifugation to a final volume of 5 ml (Vivaspin 20; 10 kDa cut-off; Sartorius), and applied to a size-exclusion chromatography column (Superdex 200, 16/60) at a flow rate of 1.5 mL/min in SEC buffer (20 mM Tris-HCl, pH 7.5, 150 mM NaCl, 1 mM TCEP). The fractions corresponding to the expected elution volumes for the purified proteins were collected, flash-frozen in liquid nitrogen, and stored at -80 °C.

### Stopped-flow experiments and fluorescence spectroscopy

All measurements were conducted using a Hi-Tech Scientific SF-61 single mixing stopped-flow system at 20 °C in SEC buffer, employing 290 nm LEDs for tryptophan fluorescence excitation. Tryptophan fluorescence was monitored through a WG 320 filter. Data collection and analysis were performed with software provided by Hi-Tech and followed standard collection and analysis procedures as set out below (17). The presented transients represent an average of 10-14 consecutive shots from the stopped-flow apparatus, ensuring a noise-to-signal ratio as close as possible to 10%. Any transients exhibiting drift between shots were excluded. The quoted concentrations reflect those in the reaction chamber post-mixing, which entails a twofold dilution from pre-mixing concentrations. Standard working volumes of 1 mL were used. Stopped-flow data were fitted to a single exponential model through a least-squares curve fit using Hi-Tech software. A large excess of cobalamin was used to maintain pseudo first order conditions facilitating the calculation of the observed binding rate constant (*k_obs_*) from the exponential change in fluorescence. Plots of *k_obs_* vs [cobalamin] yielded a straight-line plot (*k_obs_* = *k_on_* [cobalamin] + *k_off_*) from which *k_on_* was estimated from the slope of the plot. In principle, *k_off_* (and hence *K_d_* = *k_off_/k_on_*) can be estimated from the intercept value but *K_d_* was too tight, and the intercept value was not accurately defined. In some instances, the *k_off_* value could be estimated from a displacement experiment in which cobalamin was chased from its complex with the protein by addition of a large excess of a second cobalamin binding protein.

### Surface plasmon resonance

Surface plasmon resonance (SPR) experiments were performed using an LSA-XT instrument (Carterra). The sensor surface and sample deck were maintained at 20°C and 15°C, respectively. HC200M and HC30M sensor chips were utilized for Ado-Cbl and cyanocobalamin, respectively. Single-channel fluidics were primed with HBSTE buffer (10 mM HEPES, pH 7.4, 0.5 mg/mL BSA, 150 mM NaCl, 0.05% Tween 20, and 3 mM EDTA) as the running buffer. Prior to immobilization, the chip was conditioned by sequential 1-minute injections of 50 mM NaOH and 1 M NaCl.

Each protein was prepared at a concentration of 50 μg/mL in 10 mM sodium acetate, adjusted to one of four pH values (4.0, 4.5, 5.0, and 5.5). The chip surface was activated using a freshly prepared activation mixture (1:1:1 ratio of 100 mM 2-Morpholinoethanesulfonic acid (MES), pH 5.5, 100 mM sulfo-N-hydroxysuccinimide (Sulfo-NHS), and 400 mM 1-ethyl-3-(3-dimethylaminopropyl) carbodiimide hydrochloride (EDAC HCl)). Activation was carried out with a 7-minute injection, followed by direct coupling of proteins onto the chip surface using the multi-channel fluidic module via a 10-minute injection. The surface was then quenched with 0.5 M ethanolamine, pH 8.5, for 7 minutes. Following immobilization, six HBSTE buffer injections were performed to stabilize the baseline.

For analyte binding studies, cobamides were injected as a 3-fold dilution series ranging from 1.2 to 100 nM. Injection cycles consisted of a 60-second baseline stabilization, followed by a 7-minute association phase and a 20-minute dissociation phase.

SPR data for BtuF3 and BtuH1 were analyzed using a global Y-alignment method due to their relatively fast dissociation rates, whereas BtuG1, BtuG2, BtuG3, and BtuK1 were analyzed using non-regenerative kinetic modeling. Standard referencing was performed by subtracting signals from adjacent empty reference spots, followed by double referencing using a leading blank injection.

Non-regenerative kinetics were employed, ensuring that a full analyte titration series was performed without regeneration between injections. For ligands exhibiting slow dissociation (i.e., analyte binding did not return to baseline within each cycle), serial Y-alignment was applied using the baseline of the first analyte injection, followed by baseline cropping. The resulting serially Y-aligned data were fitted using a 1:1 Langmuir binding model with the T₀ float option enabled. Ligands with faster dissociation rates were analyzed using global Y-alignment and fitted to the standard 1:1 Langmuir binding model.

### Crystallization and data collection

BtuJ1 protein was concentrated to 40 mg/mL by ultrafiltration (Vivaspin 20 devices, 10 kDa cut-off, Sartorius). The initial screening for crystallization conditions was performed using commercially available sparse matrix crystallization screening kits (Molecular Dimensions) with the sitting drop vapor diffusion approach in standard 96-well crystallization plates. The total drop size was 0.4 µL, consisting of equal volumes of protein stock solution and mother liquor, with a reservoir solution volume of 100 µL. The plates were incubated at 20 °C. Further optimization of screening conditions was used the hanging drop vapor diffusion method in standard 24-well plates. The total drop size was 2.0 µL, consisting of equal volumes of protein stock solution and mother liquor, with a reservoir solution volume of 1.0 mL. The mother liquor composition was determined based on results from the sparse matrix screening experiments and was as follows: trisodium citrate: 0.05/0.1/0.2/0.3 M, pH 3.5; PEG 6000: 15/18/21/24/29/33% (w/v). The plates were stored at 20°C. For vitrification, the cryoprotectant consisting of the corresponding mother liquor solutions containing 20% glycerol (v/v) were used. Prior to X-ray data collection, the mounted crystals were stored in standard Uni-pucks (Molecular Dimensions) in liquid nitrogen. BtuJ2, protein crystals were taken directly from the screening plate of structure 2 screen 29 (2M ammonium sulfate, 0.1M sodium citrate pH 5.6, 0.2M potassium sodium tartrate tetrahydrate) and cryo-protected with mother liquor augmented with 20% glycerol. Diffraction data were collected at ESRF beamline ID23-2 (ESRF, Grenoble).

### X-ray structure determination

The X-ray diffraction data were processed using auto-processing pipelines at ESRF. The space group was determined with POINTLESS (23) and data merged using AIMLESS (24). The preliminary model was built using the AlphaFold MR workflow in the CCP4i suite and improved using ModelCraft (25), followed by rounds of manual building in Coot and refinement with REFMAC (26). MolProbity (27) was used to validate the protein geometry, and PyMOL used for the visualization of the protein structure. The coordinates and structure factors of BtuJ1 and BtuJ2 in complex with cyanocobalamin were deposited in the Protein Data Bank with identifiers 9FCT and 9I2L, respectively.

### BEV preparation and proteomic analysis

*B. thetaiotaomicron* VPI-5482 cells were grown overnight from frozen stocks in 5 mL BDMr6 in triplicate. 200 µL of these cultures were used to inoculate 5 mL BDMr6 with either 400 µM L-methionine or 1 µM cyanocobalamin and grown for 5 h. 4 mL of these cultures were then transferred into 500 mL BDMr6 with either 400 µM L-methionine or 1 µM cyanocobalamin and grown for 14 h. BEV purification and proteomic analysis was carried out as described below.

To lyse the cells/BEVs, sodium dodecyl sulphate (SDS) was added to 2% followed by boiling and vortexing. Proteins were precipitated with methanol/chloroform and the protein pellets were resuspended in 200 µl of 2.5% sodium deoxycholate (SDC) in 0.2 M EPPS-buffer pH 8, and vortexed under heating. Protein concentration was estimated using a BCA assay and 100 µg of protein per sample was treated with dithiothreitol and iodoacetamide to alkylate cysteine residues and digested with trypsin in the SDC buffer according to standard procedures. After the digest, the SDC was precipitated by adjusting to 0.2% trifluoroacetic acid (TFA), and the clear supernatant subjected to C18 SPE (Reprosil, Dr. Maisch GmbH). Peptide concentration was further estimated by running an aliquot of the digests on LCMS. Isobaric labelling was performed using a TMT™ 6plex kit (Lot VF291489, ThermoFisher Scientific) according to the manufacturer’s instructions with slight modifications; approx. 100 µg of the dried peptides were dissolved in 90 µl of 0.2 M EPPS buffer/10% acetonitrile, and 250 µg TMT reagent dissolved in 22 µl of acetonitrile was added. Samples were assigned to the TMT channels in the following order: Experiment qib43rj (BEV samples): channels 126, 127 and 128 assigned to condition M, 129, 130 and 131 to condition B; Experiment qib48rj (cell samples): channels 126, 128 and 130 assigned to condition M; channels 127, 129 and 131 to condition B.

After 2 h incubation, aliquots of 1 µL from each sample were combined in 400 µL 0.2% TFA, desalted, and analysed on the mass spectrometer (same method as for TMT, see below, but without RTS) to check labelling efficiency and estimate total sample abundances. The main sample aliquots were quenched by adding 8 µL of 5% hydroxylamine and then combined to roughly level abundances and desalted using a C18 Sep-Pak cartridge (200 mg, Waters). The eluted peptides were dissolved in 500 µL of 25 mM NH_4_HCO_3_ and fractionated by high pH reversed phase HPLC. Using an ACQUITY Arc Bio System (Waters), the samples were loaded to an XBridge® 5 µm BEH C18 130 Å column (250 x 4.6 mm, Waters). Fractionation was performed with the following gradient of solvents A (water), B (acetonitrile), and C (25 mM NH_4_HCO_3_ in water) at a flow rate of 1 mL min^-1^: solvent C was kept at 10% throughout the gradient; solvent B: 0-5 min: 5%, 5-10 min: 5-10%, 10-80 min: 10-45%, 80-90 min: 45-80%, followed by 5 min at 80% B and re-equilibration to 5% for 24 min. Fractions of 1 ml were collected and concatenated by combining fractions of similar peptide concentration to produce 16 final fractions for experiment qib43rj (BEV samples) and 21 final fractions for experiment qib48rj (cell samples) for MS analysis. Aliquots were analyzed by nanoLC-MS/MS on an Orbitrap Eclipse™ Tribrid™ mass spectrometer coupled to an UltiMate^®^ 3000 RSLCnano LC system (Thermo Fisher Scientific). The samples were loaded onto a trap cartridge (Pepmap Neo, C18, 5um, 0.3x5mm, Thermo) with 0.1% TFA at 15 µL min^-1^ for 3 min. The trap column was then switched in-line with the analytical column (nanoEase M/Z column, HSS C18 T3, 1.8 µm, 100 Å, 250 mm x 0.75 µm, Waters) for separation using the following gradient of solvents A (water, 0.1% formic acid) and B (80% acetonitrile, 0.1% formic acid) at a flow rate of 0.2 µL min^-1^ : 0-3 min 3% B (parallel to trapping); 3-10 min linear increase B to 8 %; 10-105 min increase B to 50%; 105-113 min linear increase B to 99 %; keeping at 99% B for 3 min and re-equilibration to 3% B. Data were acquired with the following parameters in positive ion mode: MS1/OT: resolution 120K, profile mode, mass range *m/z* 400-1800, AGC target 4e^5^, max inject time 50 ms; MS2/IT: data dependent analysis with the following parameters: 3 s cycle time Rapid mode, centroid mode, quadrupole isolation window 0.7 Da, charge states 2-5, threshold 1.9e^4^, CID CE = 35, AGC target 1e4, max. inject time 70 ms, dynamic exclusion 1 count for 15 s mass tolerance of 7 ppm; MS3 synchronous precursor selection (SPS): 10 SPS precursors, isolation window 0.7 Da, HCD fragmentation with CE=65, Orbitrap Turbo TMT and TMTpro resolution 30k, AGC target 1e5, max inject time 105 ms, Real Time Search (RTS): protein database *Bacteroides thetaiotaomicron* (uniprot.org, March 2022, 7482 entries), enzyme trypsin, 1 missed cleavage, oxidation (M) as variable, carbamidomethyl (C) and TMT as fixed modifications, precursor tolerance 6 ppm, Xcorr = 1, dCn = 0.05.

The acquired raw data were processed and quantified in Proteome Discoverer 3.0 (Thermo) using the incorporated search engines Comet and CHIMERYS (MSAID, Munich, Germany). The processing workflow for both engines included recalibration of MS1 spectra (RC), reporter ion quantification by most confident centroid (20 ppm) and a search on the *Bacteroides thetaiotaomicron* protein database and a common contaminants database. For CHIMERYS the Top N Peak Filter was used with 20 peaks per 100 Da and the inferys_2.1_fragmentation prediction model was used with fragment tolerance 0.5 Da, enzyme trypsin with 1 missed cleavage, variable modification oxidation (M), fixed modifications carbamidomethyl (C) and TMT6plex on N-terminus and K. For Comet the version 2019.01 rev0 parameter file was used with default settings except precursor tolerance set to 6 ppm and trypsin missed cleavages set to 1. Modifications were the same as for CHIMERYS.

The consensus workflow included the following parameters: assigning the 4 replicates/channels as described above per condition, only unique peptides (protein groups) for quantification, intensity-based abundance, TMT channel correction values applied (VF291489), co-isolation/SPS matches thresholds 50%/70%, normalised CHIMERYS Coefficient Threshold 0.8, normalisation on total peptide abundances, protein abundance-based ratio calculation, missing values imputation by low abundance resampling, hypothesis testing by t-test (background based), adjusted p-value calculation by BH-method.

### Bacteroides growth assay

For single gene rescue experiments, *B. thetaiotaomicron* VPI-5482 *Δtdk Δlocus1 Δlocus2 Δlocus3* (6) strain was transformed with single gene expression plasmids (pBATH.68-83; Supplementary Table 3). Single colonies were selected and grown in 15 mL BHIH containing 400 µM L-methionine, 250 nM cyanocobalamin and 25 µg/mL erythromycin overnight. The cells were separated using centrifugation at 3,000 g for 20 min and the supernatant was filtered via a 0.22 µm PES filter. The BEVs were then separated and concentrated using a VivaSpin6 100 kDa PES centrifugal filter followed by 5 washes with 5.5 mL PBS, concentrating to the minimal volume between washes. The final concentrated solution was resuspended in 15 mL BDMr6, containing no L-methionine, and filter sterilized using a 0.22 µm PES filter. 1 mL of the resulting BEV containing media in 24 well plate (bio-one cellstar; Greiner) was inoculated with 50 µL *B. thetaiotaomicron* VPI-5482 *Δtdk* pre-cultured in BDMr6 sealed with adhesive gas permeable seals to limit evaporation. A negative control was included for each BEV containing BDMr6 condition and a positive control containing 200 µM L-methionine was used. Plates were incubated overnight, and cell growth was evaluated by OD_600_. Note that *B. thetaiotaomicron* VPI-5482 must be pre-cultured in BDMr6 with L-methionine and lacking any source of cobalamin for several generations for the assay to work. This ensures that any stored cobalamin is depleted prior to the bioassay.

For *ΔbtuJ1* and *ΔbtuJ2* experiments, wild-type *B. thetaiotaomicron* VPI-5482 was used as the strain to generate knockouts and as the bioassay test strain. *B. thetaiotaomicron* VPI-5482 Δ1234 strain, specifically lacking all of the uptake operons was also constructed as *B. thetaiotaomicron* VPI-5482 *Δtdk Δlocus1 Δlocus2 Δlocus3* strain contains additional gene knockouts found between operons 3 and 4. Two mutant colonies selected for each strain and cultured in 15 mL BDMr6 containing 400 µM L-methionine to mid-log phase. Further assay steps carried out as above, except cyanocobalamin added to 250 nM to the filtered supernatant prior to BEV preparation. Assays carried out in 1 mL in 48 well plates (bio-one cellstar; Greiner) sealed with adhesive gas permeable seals to limit evaporation. Plates were incubated overnight and OD_600_ readings were taken indicating cell growth. Each BEV preparation was tested in four replicates.

For cobalamin uptake operon mutant analysis, two separate colonies of the different mutant strains were inoculated in duplicate (four replicates) in 5 mL BDMr6 and grown overnight. 50 µL of was then used to inoculate 1 mL BDMr6 in 48 well plates (as above), where L-methionine was replaced with different amounts of cyanocobalamin or BEVs prepared from wild-type *B. thetaiotaomicron* VPI-5482 as in the *ΔbtuJ1* and *ΔbtuJ2* experiment. Cells growth evaluated as above.

BEV release overtime experiments carried out based on previously described method with slight modifications (10). Two colonies for each strain were selected and used to inoculate 5 mL BDMr6 (200 µM L-methionine) and grown overnight. 1 mL was then sub-cultured into 300 mL BDMr6 (200 µM L-methionine) and grown for 16 h. Cultured were then centrifuged in sterile bottles pre-equilibrated in anaerobic cabinet at 6,600 g at 21 °C for 10 min. Pelleted cells resuspended in BDMr6 and used to inoculate 300 mL BDMr6 to starting OD_600_ of 0.25. Lipid concentration evaluated using FM4-64 dye based assay (13).

*B. fragilis* NCTC 9343 BEV utilization was evaluated using the bioassay as described for *ΔbtuJ1* and *ΔbtuJ2* experiments.

## Supporting information

Supplemental information is provided

## Funding

This work was supported by the Biotechnology and Biological Sciences Research Council (BBSRC) under Grant BB/R012490/1; BBSRC under Grant BB/CCG1860/1; BBSRC under Grant BB/S002197/1; and Royal Society under Grant INF/R2/180062.

## CRediT Author Contribution

RJ: Conceptualization; Methodology; Formal analysis; Investigation; Writing - Original Draft; Writing - Review & Editing. RU: Conceptualization; Methodology; Formal analysis; Investigation; Writing - Original Draft. CC: Methodology; Formal analysis; Writing - Original Draft. MB: Methodology; Formal analysis. ED: Methodology; Supervision. CM: Methodology; Investigation; Writing - Original Draft. GS: Methodology; Formal analysis. BK: Methodology, Formal analysis. SRC: Conceptualization; Writing - Review & Editing; Supervision; Funding acquisition. MG: Methodology; Supervision; Writing - Original Draft. RWP: Conceptualization; Writing - Original Draft; Writing - Review & Editing; Supervision; Funding acquisition. MW: Conceptualization; Writing - Original Draft; Writing - Review & Editing; Supervision; Funding acquisition.

## Data availability statement

The authors confirm that data supporting the findings of this study are available within the article or are available from the corresponding author on reasonable request.

## Competing Interests

The authors declare that there are no competing interests associated with the manuscript.

## Notes

https://www.rcsb.org/search?request=%7B%22query%22%3A%7B%22type%22%3A%22group%22%2C%22nodes%22%3A%5B%7B%22type%22%3A%22group%22%2C%22nodes%22%3A%5B%7B%22type%22%3A%22group%22%2C%22nodes%22%3A%5B%7B%22type%22%3A%22terminal%22%2C%22service%22%3A%22full_text%22%2C%22parameters%22%3A%7B%22value%22%3A%22BtuJ1%22%7D%7D%5D%2C%22logical_operator%22%3A%22and%22%7D%5D%2C%22logical_operator%22%3A%22and%22%2C%22label%22%3A%22full_text%22%7D%5D%2C%22logical_operator%22%3A%22and%22%7D%2C%22return_type%22%3A%22entry%22%2C%22request_options%22%3A%7B%22paginate%22%3A%7B%22start%22%3A0%2C%22rows%22%3A25%7D%2C%22results_content_type%22%3A%5B%22experimental%22%5D%2C%22sort%22%3A%5B%7B%22sort_by%22%3A%22score%22%2C%22direction%22%3A%22desc%22%7D%5D%2C%22scoring_strategy%22%3A%22combined%22%7D%2C%22request_info%22%3A%7B%22query_id%22%3A%225bd40b7d1ff43dbae1df17ba0f70e1c1%22%7D%7D

https://www.rcsb.org/search?request=%7B%22query%22%3A%7B%22type%22%3A%22group%22%2C%22nodes%22%3A%5B%7B%22type%22%3A%22group%22%2C%22nodes%22%3A%5B%7B%22type%22%3A%22group%22%2C%22nodes%22%3A%5B%7B%22type%22%3A%22terminal%22%2C%22service%22%3A%22full_text%22%2C%22parameters%22%3A%7B%22value%22%3A%22BtuJ2%22%7D%7D%5D%2C%22logical_operator%22%3A%22and%22%7D%5D%2C%22logical_operator%22%3A%22and%22%2C%22label%22%3A%22full_text%22%7D%5D%2C%22logical_operator%22%3A%22and%22%7D%2C%22return_type%22%3A%22entry%22%2C%22request_options%22%3A%7B%22paginate%22%3A%7B%22start%22%3A0%2C%22rows%22%3A25%7D%2C%22results_content_type%22%3A%5B%22experimental%22%5D%2C%22sort%22%3A%5B%7B%22sort_by%22%3A%22score%22%2C%22direction%22%3A%22desc%22%7D%5D%2C%22scoring_strategy%22%3A%22combined%22%7D%2C%22request_info%22%3A%7B%22query_id%22%3A%2230188c190db6948cafd0c116bc801e93%22%7D%7D

### Competing Interest Statement

The authors have declared no competing interest.

