## Supplemental information is provided for "Extracellular Vesicle-Linked Vitamin B_12_ Acquisition via Novel Binding Proteins in *Bacteroides thetaiotaomicron*"

### Supporting Information for
#### Extracellular Vesicle-Linked Vitamin $B_{12}$ Acquisition via Novel Binding Proteins in *Bacteroides thetaiotaomicron*

Rokas Juodeikis, Robert Ulrich, Charlea Clarke, Michal Banasik, Evelyne Deery, Gerhard Saalbach, Bernhard Krautler, Simon R. Carding, Michael A. Geeves, Richard W. Pickersgill, and Martin J. Warren

**This PDF file includes:**

Figures S1 to S4
Tables S1 to S3
Legends for Datasets S1 to S3

**Other supporting materials for this manuscript include the following:**

Datasets S1 to S3

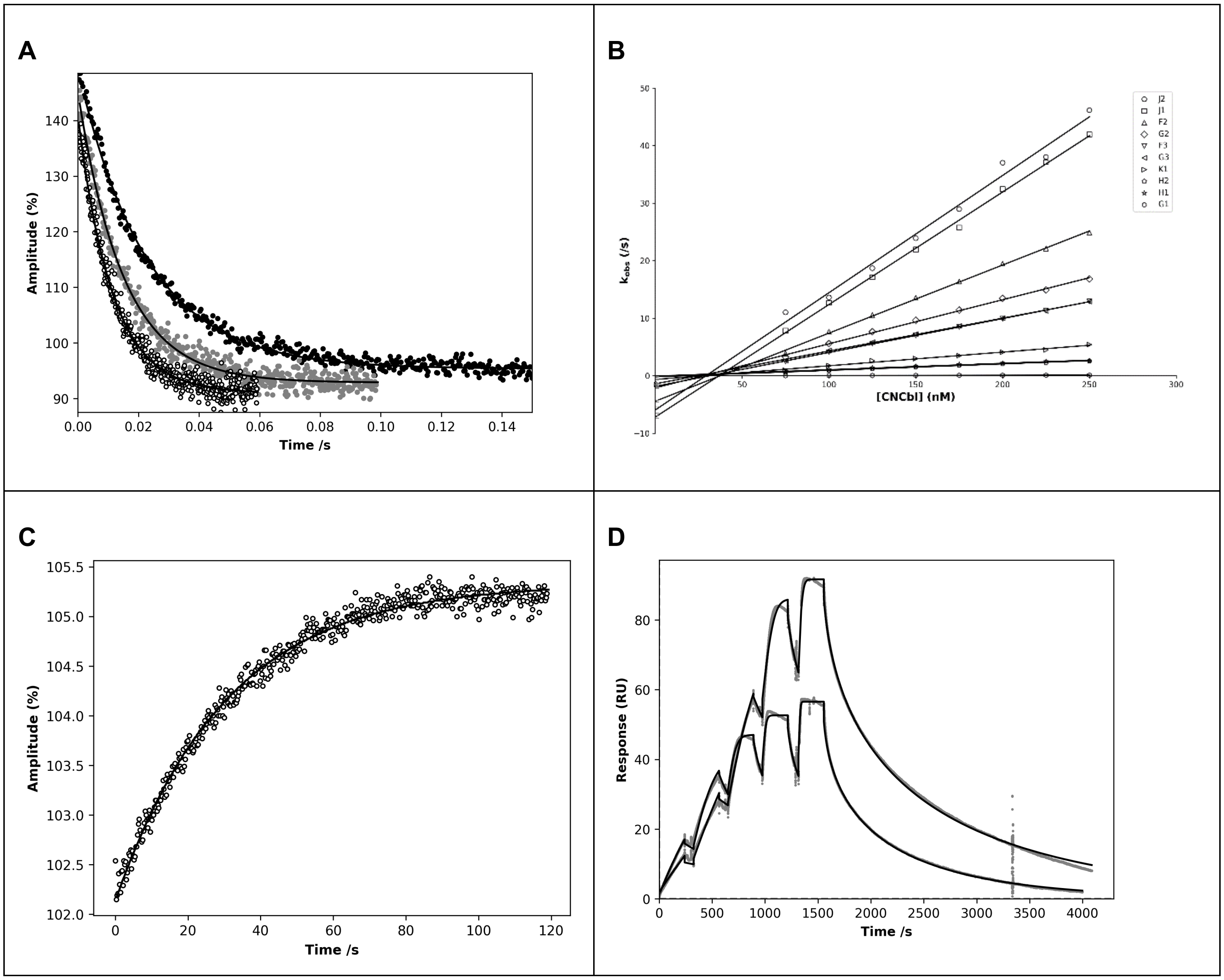


**Fig. S1. The binding kinetics of BtuJ1 and other B12 binding proteins. A**) A representative dataset to establish $k_{on}$. The change in tryptophan fluorescence upon BtuJ1 binding to CN-Cbl using 25nM BtuJ1. Black, gray and white circles are 75 nM, 150 nM and 225 nM CN-Cbl, respectively; from which $k_{on}$ can be determined to be 0.20 /nM/s +/- 0.0036 /nM/s.

**B**) Linear regression curves. $k_{obs}$ for 10 proteins binding CN-Cbl was plotted against a series of CN-Cbl concentrations. All reactions contain 25 nM protein. A remarkable range of off rates can be seen with BtuJ2 forming a complex with CN-Cbl the fastest where $k_{on}$ = 0.2 /nM/s and BtuG1 was the exception with a very slow rate of forming a complex with CN-Cbl where $k_{on}$ = 0.0004 nM-1s-1. The key is ordered by decreasing $k_{on}$ value.

**C**) Measuring $k_{off}$. An example of the change in tryptophan fluorescence upon BtuJ2 displacing a complex of BtuJ1 with CN-Cbl from which Koff can be established to be 0.032/s.

**D**) A representative SPR sensogram displaying the binding interaction of BtuJ1 and CN-Cbl. The two traces represent duplicate data. Values for $k_{on}$ and $k_{off}$ were calculated from the mean of two repeats. SE values are displayed in parentheses. $k_{on}$ – 0.014/nM/s (+/- 0.00020/nM/s). $k_{off}$ – 0.010/s (+/- 0.00010). $K_d$ = 0.70 nM (+/- 0.010 nM). $k_{on}$ = 0.2/nM/s and BtuG1 forming a complex with CN-Cbl the slowest where $k_{on}$ = 0.0004 /nM/s. The key is ordered by decreasing $k_{on}$ value.

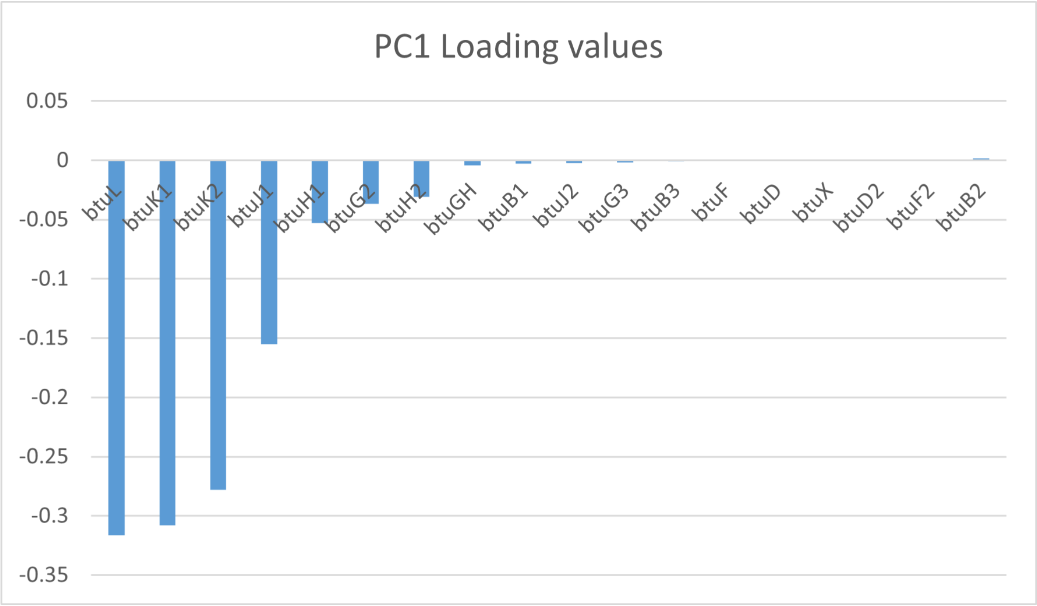


**Fig. S2. PCA analysis PC1 loading values for identified cobamide uptake operon proteins.** PCA analysis shows that PC1 negative loading values are indicative of proteins enriched in non-lytic BEVs released during growth phase. Only specific cobamide uptake operon proteins shows such values, suggesting specific enrichment in non-lytic BEVs.

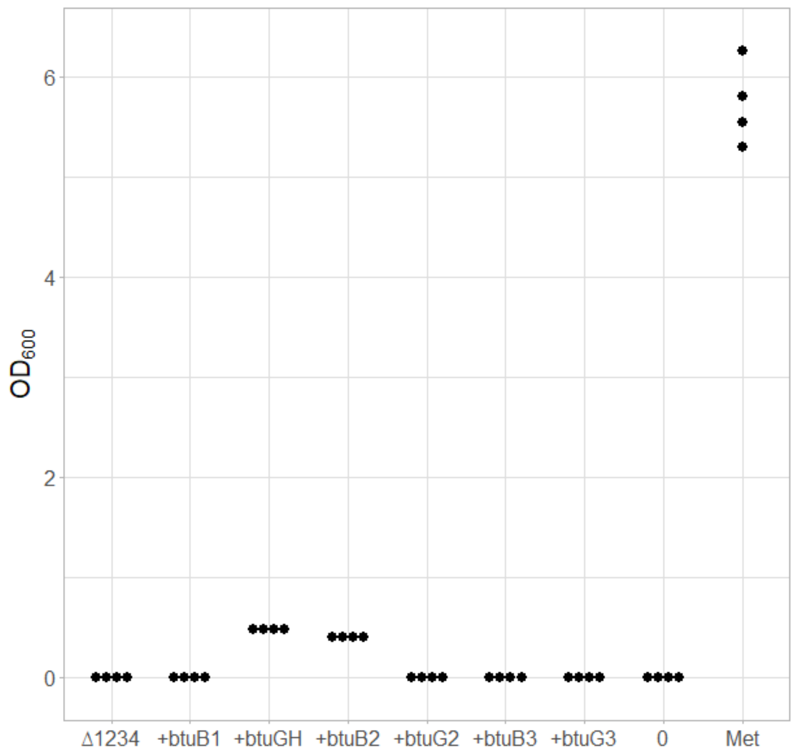


**Fig. S3.** ***B. thetaiotaomicron*** **cobalamin bioassay results showing that BtuG and BtuB are not responsible for cobalamin delivery on BEVs.** Bioassay carried out in four biological replicates with one biological BEV replicate.

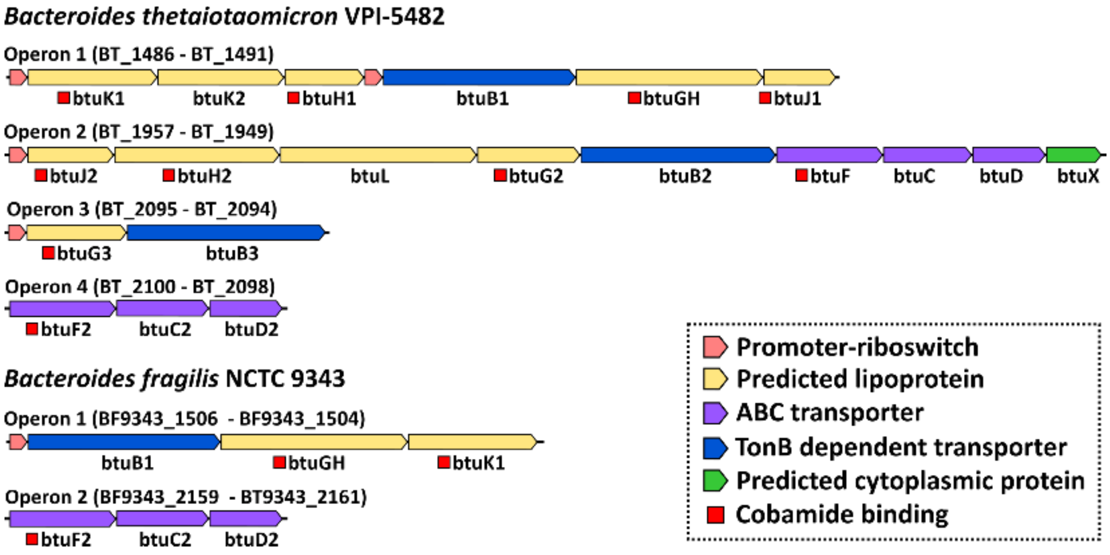


**Fig. S4. *B. thetaiotaomicron* and *B. fragilis* NCTC 9343 cobalamin uptake operons.** *B. fragilis* has a significantly smaller number of genes within the cobalamin uptake operons. Notably, no copies of *btuJ* are present.

**Table S1. X-ray data collection and refinement statistics for BtuJ1 and BtuJ2**

| **Protein (PDB identifier)** | **BtuJ1** | **BtuJ2** |
|---|---|---|
| **Data collection** | | |
| Source | ESRF (Grenoble) | DLS (Oxford) |
| Wavelength | 0.873128 | 0.95370 |
| Space group | C2 | $P2_12_12$ |
| Cell parameters (Å) | 148.9, 51.4,108.6, β=131.8° | 83.75, 97.98, 70.47 |
| Resolution (higher shell) | 81.1 - 1.60 (1.70-1.60) | 70.5 – 2.44 (2.44-2.66) |
| $R_{pim}$ (all reflections) | 0.120 (0.841) | 0.192 (0.661) |
| Mean I/sd(I) | 4.8 (1.6) | 6.2 (1.4) |
| Completeness (%) | 90.0 (42.2) | 88.8 (64.5) |
| Multiplicity | 4.4 (4.4) | 12.6 (13.1) |
| Wilson B-factor ($Å^2$) | 13.6 | 32.6 |
| **Refinement** | | |
| Number of reflections (working/test) | 62828/3429 | 14879/790 |
| $R_{work}$ | 0.189 | 0.196 |
| $R_{free}$ | 0.214 | 0.257 |
| Protein atoms | 3860 | 4269 |
| Number of water atoms modeled | 496 | 36 |
| Ligand (atoms) | CN-Cbl (B12) | Cn-Cbl (B12) |
| RMSD Bond lengths (Å) | 0.013 | 0.007 |
| RMSD Bond angles (˚) | 2.332 | 2.022 |
| Ramachandran preferred (%) | 97.2 | 94.3 |
| Ramachandran allowed (%) | 2.8 | 4.1 |
| Ramachandran outliers (%) | 0 | 1.6 |
| MolProbity score | 1.04 | 2.20 |

**Table S2. Primers used in this study**

| PrimerID | Sequence |
|---|---|
| 397FwPromRBS1486bgl | GTCAGATCTCATGTAGTTCCAATATCAGTTCCAAC |
| 398RvPromRBS1486nde | GATCATATGTTCTCTGAATCAGAGTGATGAATG |
| 399FwPromRBS1489bgl | GTCAGATCTGTATCCGAAGACGCTAATGGCAAC |
| 400RvPromRBS1489nde | CATATGTTTCTTTCAGTCTGTTCATAATGTAAAAAGATG |
| 401FwPromRBS1957bgl | GTCAGATCTGCTGGAAACTCTTCAGAAAGTG |
| 402RvPromRBS1957nde | GATCATATGTCACTCATTTTAATGTAAAACATATTATTAACAAAC |
| 403FwPromRBS2095bgl | GTCAGATCTGTATTATATCTTATAATGATACCCCTTTGAAAG |
| 404RvPromRBS2095nde | GATCATATGTAAAAGCGAGAAAAGGTTATTAACTAACTTG |
| 405FwPromRBS2098bgl | GTCAGATCTGAAACAGAAATCAAGCTTTCTTC |
| 406RvPromRBS2098nde | GCTCATATGGATGTTTTTATCTGAGGACAAAA |
| BT1486_NdeI_FOR | GCACATATGTGCTCGAAAGACGATTGTG |
| BT1486_SpeI_REV | TCGACTAGTTAGTTAAAAATGAATGTAGCCGG |
| BT1487_NdeI_FOR | CGACATATGTGTAACGACGACGATTG |
| BT1487_SpeI_REV | CGAACTAGTTAGTTATTGAATAATATTTGTGAAGGAAC |
| BT1488_NdeI_FOR | GCACATATGTGCAACAAAGATGAGGAAG |
| BT1488_SpeI_REV | GCAACTAGTTAAAAAGCTTTGAATCCAACCACCTCG |
| BtBtuGH_NdeI | CATCATATGTGCGATGATCTGGAAGATAAG |
| BtBtuGH_SpeI | CATACTAGTTAGTTCTCGAAATGTAAATC |
| BT1491_NdeI_FOR | GCACATATGTGTAGTGATGATGATGAG |
| BT1491_SpeI_REV | GCAACTAGTTATTTTTCGATGAGGGTGATACC |
| BT1957_NdeI_FOR | GCACATATGTGTAGCTCGGACGATGAC |
| BT1957_SpeI_REV | GCAACTAGTTATTGTATGGTGATATCATC |
| BT1956_NdeI_FOR | GCACATATGTGCAACAAGGATGAAGTC |
| BT1956_SpeI_REV | GCAACTAGTTATTTGGTTAGATCCTC |
| BT1955_NdeI_FOR | GCACATATGTGTGACAAAAACGATG |
| BT1955_SpeI_REV | GCAACTAGTTAATCCTGCTGCGTATATTG |
| BtBtuG2_Nde | CATACATATGGAAGATTTCTCTGTATCG |
| BtBtuG2_Spe | ATGACTAGTTATTTCCAGCAGAAAGCTCC |
| BT1952_NdeI_FOR | GCACATATGTGCCACAACAAAAGCTC |
| BT1952_SpeI_REV | GCAACTAGTCTATTTCAGTTGCTTG |
| BtBtuG3_Nde | CATCATATGTGTATGAAATGGGATTATG |
| BtBtuG3_Spe | AGTACTAGTTACTTCCAACAAAATGC |
| BT2098_NdeI_FOR | GCACATATGTGCGTATACAATAAAAAAACTTCTTTGG |
| BT2098_SpeI_REV | GCAACTAGTCATTCTAGATGTCTG |
| 274FwBT1486nde | GATCATATGAATTGTAAAAAGCTATTCAAAACGTTATTATTTA |
| 275RvBT1486spe | GATACTAGTTTAGTTAAAAATGAATGTAGCCGGGAAATAATAG |
| 276FwBT1487nde | GATCATATGAATAAACTATATACCACTTTATTAATAGCCTG |
| 277RvBT1487spe | GATACTAGTTTAGTTATTGAATAATATTTGTGAAGGAACGCTC |
| 278FwBT1488nde | GATCATATGAAAAGATATTGGTATCTGATGGCTATAG |
| 279RvBT1488spe | GATACTAGTTTAAAAAGCTTTGAATCCAACCACCTC |
| 280FwBT1489nde | GATCATATGAGAAGGAATACTTTTATTAAAAAGATGAGCGTAC |

| | |
|---|---|
| 281RvBT1489spe | CTGACTAGTCTAATACCTGACTCCTATCGTCACTC |
| 282FwBT1490nde | GTACATATGCAGAAAGGTCTTTTATATAATATGTTG |
| 283RvBT1490spe | GATACTAGTTTAGTTCTCGAAATGTAAATCCTCTAC |
| 284FwBT1491nde | GATCATATGAAAGCAAAAATGAAAAAGTTATCTTTATTC |
| 285RvBT1491spe | GATACTAGTCTATTTATTTTTCGATGAGGGTGATAC |
| 286FwBT1957nde | GATCATATGAAAAGAAAATTACGCTTTCTGGCAG |
| 287RvBT1957spe | GATACTAGTTTATTGTATGGTGATATCATCAATACAGATATAAG |
| 288FwBT1956nde | GATCATATGCATCGTTTTCACTATTTTATTATTTCTG |
| 289RvBT1956spe | GATACTAGTTTATTTGGTTAGATCCTCAAAAGAAAATAC |
| 291FwBT1954nde | GTACATATGATTCGGGTACTCTTTTTTATCCGAATG |
| 292RvBT1954spe | GATACTAGTTTATTTCCAGCAGAAAGCTCCCGGAATG |
| 293FwBT1953nde | GTACATATGAAAAGACATCTTATTCTATTGTTCGTG |
| 294RvBT1953spe | GATACTAGTTTATCGTTTACTATTTTTGTTTTTTCCGAACTTG |
| 303FwBT2095nde | GATCATATGAAACGAATTTTACTTTCTGTTTTATTTATTGTCTTCTG |
| 304RvBT2095spe | GATACTAGTTTACTTCCAACAAAATGCTCCGGGAATAATTC |
| 305FwBT2094nde | GTACATATGAGGAGAAATATATTATTAGTGCAGTTTGTAGGAGTTC |
| 306RvBT2094spe | GATACTAGTCTATTTTTTCTTCTTGCCCCACTTGGGAG |
| 341FwUpBT1491bam | CTAGGATCCAGTCAAAGGAGATGTCCTTTGC |
| 342RvUpBT1491 | GTGATACCGTCAAGGATTAGTTCTCGAAATGTAAATCCTC |
| 343FwDownBT1491 | CATTTCGAGAACTAATCCTTGACGGTATCACCCTCATC |
| 344RvDownBT1491pst | CTACTGCAGGTATTTATTGGCGATAGCACGTAC |
| 349FwUpBT1957Bam | CTAGGATCCTGCTGGAAACTCTTCAGAAAGTG |
| 350RvUpBT1957 | CAATACAGATATAAGCGTCACTCATTTTAATGTAAAACATATTATTAAC |
| 351FwDownBT1957 | CATTAAAATGAGTGACGCTTATATCTGTATTGATGATATCACCATAC |
| 352RvDownBT1957pst | CTACTGCAGACTTTTACGGTGGCAGTGAC |
| 386FwUpBT1486-91bam | CTAGGATCCGGTATTGGAAGCAATGATAGCCTCTCC |
| 387RvUpBT1486-91 | GATACCGTCAAGCTCATGCTCTAACAATTCCTGCGTATC |
| 388FwDownBT1486-91 | GTTAGAGCATGAGCTTGACGGTATCACCCTCATCG |
| 344RvDownBT1491pst | CTACTGCAGGTATTTATTGGCGATAGCACGTAC |
| 382FwUpBT1957-49bam | CTAGGATCCATTCGAGTGTATGCCCGAATAC |
| 383RvUpBT1957-49 | GACGGTTTAACCACACAAAGCATGACAACCGTAC |
| 384FwDownBT1957-49 | ATGCTTTGTGTGGTTAAACCGTCCTGTTTTCGTTAAAC |
| 385RvDownBT1957-49pst | CTACTGCAGACCTGACTTCCATGACTTGGTATCTC |
| 389FwUpBT2094-5bam | CTAGGATCCGTTGGTAGTTGATTATGTGATCGAC |
| 390RvUpBT2094-5 | CGATAAAGAACCAGGAACAAATCTGATGAACAGAATC |
| 391FwDownBT2094-5 | GATTTGTTCCTGGTTCTTTATCGGTATAACTCCCAAG |
| 392RvDownBT2094-5pst | CTACTGCAGCATTATCTATGTTCCTTCGTATCCTGAC |
| 393FwUpBT2098-100bam | CTAGGATCCTGATTCCGCAAATTAAAGCTG |
| 394RvUpBT2098-100 | GTAAGGGAGGTGCAAGCATGACAGGCGGAAATTATC |
| 395FwDownBT2098-100 | CTGTCATGCTTGCACCTCCCTTACGAGAAAGTTTC |
| 396RvDownBT2098-100pst | CTACTGCAGAACTGGAACGCGTTCTATACAAC |

**Table S3. Plasmids used in this study**

| PlasmidID | Description |
|---|---|
| TetR-P1T_DP-GH023 | Single copy *B. thetaiotaomicron* integration vector. Obtained from addgene (Plasmid #90324). |
| pIBATH.56 | TetR-P1T_DP-GH023 *AflII/BamHI* site replaced with a synthetic DNA fragment containing a promoter (P.Bth_BT1830s); ribosome binding site (RBS.Bth_RBS7) and a codon optimized Nanoluciferase; flanked by terminators (T.BBa_B1001; T.BBa_B1007). |
| pIBATH.104 | pIBATH.56 *BglII/NdeI* site replaced by a *BglII/NdeI* DNA fragment amplified using primers 397FwPromRBS1486bgl and 398RvPromRBS1486nde corresponding to *btuK1* (BT_1486) RBS and upstream promoter (operon 1). |
| pIBATH.105 | pIBATH.56 *BglII/NdeI* site replaced by a *BglII/NdeI* DNA fragment amplified using primers 399FwPromRBS1489bgl and 400RvPromRBS1489nde corresponding to *btuB1* (BT_1489) RBS and upstream promoter (operon 1; promoter 2). |
| pIBATH.106 | pIBATH.56 *BglII/NdeI* site replaced by a *BglII/NdeI* DNA fragment amplified using primers 401FwPromRBS1957bgl and 402RvPromRBS1957nde corresponding to *btuJ2* (BT_1957) RBS and upstream promoter (operon 2). |
| pIBATH.107 | pIBATH.56 *BglII/NdeI* site replaced by a *BglII/NdeI* DNA fragment amplified using primers 403FwPromRBS2095bgl and 404RvPromRBS2095nde corresponding to *btuG3* (BT_2095) RBS and upstream promoter (operon 3). |
| pIBATH.108 | pIBATH.56 *BglII/NdeI* site replaced by a *BglII/NdeI* DNA fragment amplified using primers 405FwPromRBS2098bgl and 406RvPromRBS2098nde corresponding to *btuF2* (BT_2098) RBS and upstream promoter (operon 4). |
| pET14b_bth_BT1486 | T7 promoter driven expression plasmid for 6xHis N-terminus tagged *btuK1* (BT_1486); PCR amplified *btuK1* (primers: BT1486_NdeI_FOR; BT1486_SpeI_REV) inserted into a modified pET14b vector (novel *SpeI* site between *NdeI* and terminator) *NdeI/SpeI* site. |
| pET14b_bth_BT1487 | T7 promoter driven expression plasmid for 6xHis N-terminus tagged *btuK2* (BT_1487); PCR amplified *btuK2* (primers: BT1487_NdeI_FOR; BT1487_SpeI_REV) inserted into a modified pET14b as for pET14b_bth_BT1486. |
| pET14b_bth_BT1488 | T7 promoter driven expression plasmid for 6xHis N-terminus tagged *btuH1* (BT_1488); PCR amplified *btuH1* (primers: BT1488_NdeI_FOR; BT1488_SpeI_REV) inserted into a modified pET14b as for pET14b_bth_BT1486. |
| pET14b_bth_BT1490 | T7 promoter driven expression plasmid for 6xHis N-terminus tagged *btuGH* (BT_1490); PCR amplified *btuGH* (primers: BtBtuGH_NdeI; BtBtuGH_SpeI) inserted into a modified pET14b as for pET14b_bth_BT1486. |

| | |
|---|---|
| pET14b_bth_BT1491 | T7 promoter driven expression plasmid for 6xHis N-terminus tagged *btuJ1* (BT_1491); PCR amplified *btuJ1* (primers: BT1491_NdeI_FOR; BT1491_SpeI_REV) inserted into a modified pET14b as for pET14b_bth_BT1486. |
| pET14b_bth_BT1957 | T7 promoter driven expression plasmid for 6xHis N-terminus tagged *btuJ2* (BT_1957); PCR amplified *btuJ2* (primers: BT1957_NdeI_FOR; BT1957_SpeI_REV) inserted into a modified pET14b as for pET14b_bth_BT1486. |
| pET14b_bth_BT1956 | T7 promoter driven expression plasmid for 6xHis N-terminus tagged *btuH2* (BT_1956); PCR amplified *btuH2* (primers: BT1956_NdeI_FOR; BT1956_SpeI_REV) inserted into a modified pET14b as for pET14b_bth_BT1486. |
| pET14b_bth_BT1955 | T7 promoter driven expression plasmid for 6xHis N-terminus tagged *btuL* (BT_1955); PCR amplified *btuL* (primers: BT1955_NdeI_FOR; BT1955_SpeI_REV) inserted into a modified pET14b as for pET14b_bth_BT1486. |
| pET14b_bth_BT1954 | T7 promoter driven expression plasmid for 6xHis N-terminus tagged *btuG2* (BT_1954); PCR amplified *btuG2* (primers: BtBtuG2_Nde; BtBtuG2_Spe) inserted into a modified pET14b as for pET14b_bth_BT1486. |
| pET14b_bth_BT1952 | T7 promoter driven expression plasmid for 6xHis N-terminus tagged *btuF* (BT_1952); PCR amplified *btuF* (primers: BT1952_NdeI_FOR; BT1952_SpeI_REV) inserted into a modified pET14b as for pET14b_bth_BT1486. |
| pET14b_bth_BT2095 | T7 promoter driven expression plasmid for 6xHis N-terminus tagged *btuG3* (BT_2095); PCR amplified *btuG3* (primers: BtBtuG3_Nde; BtBtuG3_Spe) inserted into a modified pET14b as for pET14b_bth_BT1486. |
| pET14b_bth_BT2098 | T7 promoter driven expression plasmid for 6xHis N-terminus tagged *btuF2* (BT_2098); PCR amplified *btuF2* (primers: BT2098_NdeI_FOR; BT2098_SpeI_REV) inserted into a modified pET14b as for pET14b_bth_BT1486. |
| pGH117 | Vector backbone used to construct single gene expression plasmids. |
| pBATH.03 | pGH117 *Pci/BamHI* site replaced with a synthetic DNA fragment containing a synthetic promoter (P.Bth_BT1830s) and ribosome binding site (RBS.Bth_RBS7) followed by an *NdeI/SpeI* cloning site containing an ORF (not relevant to this study) with artificial bidirectional terminators (BBa_B1001; BBa_B1007) flanking the operon yielding pBATH.03. |
| pBATH.68 | pBATH.03 *NdeI/SpeI* site replaced by a *NdeI/SpeI* DNA fragment amplified using primers 274FwBT1486nde and 275RvBT1486spe corresponding to *btuK1* (BT_1486). |
| pBATH.69 | pBATH.03 *NdeI/SpeI* site replaced by a *NdeI/SpeI* DNA fragment amplified using primers 276FwBT1487nde and 277RvBT1487spe corresponding to *btuK2* (BT_1487). |

| | |
|---|---|
| pBATH.70 | pBATH.03 *NdeI/SpeI* site replaced by a *NdeI/SpeI* DNA fragment amplified using primers 278FwBT1488nde and 279RvBT1488spe corresponding to *btuH1* (BT_1488). |
| pBATH.71 | pBATH.03 *NdeI/SpeI* site replaced by a *NdeI/SpeI* DNA fragment amplified using primers 280FwBT1489nde and 281RvBT1489spe corresponding to *btuB1* (BT_1489). |
| pBATH.72 | pBATH.03 *NdeI/SpeI* site replaced by a *NdeI/SpeI* DNA fragment amplified using primers 282FwBT1490nde and 283RvBT1490spe corresponding to *btuGH* (BT_1490). |
| pBATH.73 | pBATH.03 *NdeI/SpeI* site replaced by a *NdeI/SpeI* DNA fragment amplified using primers 284FwBT1491nde and 285RvBT1491spe corresponding to *btuJ1* (BT_1491). |
| pBATH.74 | pBATH.03 *NdeI/SpeI* site replaced by a *NdeI/SpeI* DNA fragment amplified using primers 286FwBT1957nde and 287RvBT1957spe corresponding to *btuJ2* (BT_1957). |
| pBATH.75 | pBATH.03 *NdeI/SpeI* site replaced by a *NdeI/SpeI* DNA fragment amplified using primers 288FwBT1956nde and 289RvBT1956spe corresponding to *btuH2* (BT_1956). |
| pBATH.77 | pBATH.03 *NdeI/SpeI* site replaced by a *NdeI/SpeI* DNA fragment amplified using primers 291FwBT1954nde and 292RvBT1954spe corresponding to *btuG2* (BT_1954). |
| pBATH.78 | pBATH.03 *NdeI/SpeI* site replaced by a *NdeI/SpeI* DNA fragment amplified using primers 293FwBT1953nde and 294RvBT1953spe corresponding to *btuB2* (BT_1953). |
| pBATH.83 | pBATH.03 *NdeI/SpeI* site replaced by a *NdeI/SpeI* DNA fragment amplified using primers 303FwBT2095nde and 304RvBT2095spe corresponding to *btuG3* (BT_2095). |
| pBATH.84 | pBATH.03 *NdeI/SpeI* site replaced by a *NdeI/SpeI* DNA fragment amplified using primers 305FwBT2094nde and 306RvBT2094spe corresponding to *btuB3* (BT_2094). |
| pLGB13 | Vector for generating clean *Bacteroides* knockout strains. Obtained from addgene (Plasmid #126618). |
| pKO.026 | pLGB13 *BamHI/PstI* site replaced with fragment of *btuJ1* flanking DNA sequences generated by overlap extension PCR; Up fragment (Primers: 341FwUpBT1491bam/342RvUpBT1491); Down fragment (Primers: 343FwDownBT1491/344RvDownBT1491pst). Used to generate Δ*btuJ1* knockout. |
| pKO.028 | pLGB13 *BamHI/PstI* site replaced with fragment of *btuJ2* flanking DNA sequences generated by overlap extension PCR; Up fragment (Primers: 349FwUpBT1957Bam/350RvUpBT1957); Down fragment (Primers: 351FwDownBT1957/352RvDownBT1957pst). Used to generate Δ*btuJ2* knockout. |
| pKO.039 | pLGB13 *BamHI/PstI* site replaced with fragment of operon 1 flanking DNA sequences generated by overlap extension PCR; Up fragment (Primers: |

| | |
|---|---|
| | 386FwUpBT1486-91bam/387RvUpBT1486-91); Down fragment (Primers: 388FwDownBT1486-91/344RvDownBT1491pst). Used to generate Δ1 (operon 1) knockout. |
| pKO.038 | pLGB13 *BamHI/PstI* site replaced with fragment of operon 2 flanking DNA sequences generated by overlap extension PCR; Up fragment (Primers: 382FwUpBT1957-49bam/383RvUpBT1957-49); Down fragment (Primers: 384FwDownBT1957-49/385RvDownBT1957-49pst). Used to generate Δ2 (operon 2) knockout. |
| pKO.040 | pLGB13 *BamHI/PstI* site replaced with fragment of operon 3 flanking DNA sequences generated by overlap extension PCR; Up fragment (Primers: 389FwUpBT2094-5bam/390RvUpBT2094-5); Down fragment (Primers: 391FwDownBT2094-5/392RvDownBT2094-5pst). Used to generate Δ3 (operon 3) knockout. |
| pKO.041 | pLGB13 *BamHI/PstI* site replaced with fragment of operon 4 flanking DNA sequences generated by overlap extension PCR; Up fragment (Primers: 393FwUpBT2098-100bam/394RvUpBT2098-100); Down fragment (Primers: 395FwDownBT2098-100/396RvDownBT2098-100pst). Used to generate Δ4 (operon 4) knockout. |
| pKO.030 | pLGB13 *BamHI/PstI* site replaced with fragment of BtuJ2/BtuH2/BtuL flanking DNA sequences generated by overlap extension PCR; Up fragment (Primers: 349FwUpBT1957bam/357RvUpBT1957); Down fragment (Primers: 358FwDownBT1955/348RvDownBT1955pst). Used to generate ΔJ2.H2.L knockout. |

**Dataset S1 (separate file).** Annotated operon and plasmid sequences and maps including primers and cloning information (.gz).

**Dataset S2 (separate file).** Comparative proteomics dataset of *B. thetaiotaomicron* cells grown with or without (400 μM L-methionine) cobalamin.

**Dataset S3 (separate file).** Comparative proteomics dataset of BEVs purified from *B. thetaiotaomicron* cultures grown with or without (400 μM L-methionine) cobalamin.
